# A general role for MIA3/TANGO1 in secretory pathway organization and function

**DOI:** 10.1101/2021.02.24.432632

**Authors:** Janine McCaughey, Nicola L. Stevenson, Judith M. Mantell, Chris R. Neal, Alex Paterson, Kate Heesom, David J. Stephens

## Abstract

Complex machinery is required to drive secretory cargo export from the endoplasmic reticulum, an essential process in eukaryotic cells. In vertebrates, the Mia3 gene encodes two major forms of Transport ANd Golgi Organization Protein 1 (TANGO1S and TANGO1L). Here, using genome engineering of human cells, light microscopy, secretion assays, genomics, and proteomics we show that disruption of the longer form, TANGO1L, results in relatively minor defects in secretory pathway organization and function including limited impacts on procollagen secretion. In contrast, loss of both long and short forms results in major defects in cell organization and secretion. These include a failure to maintain the localization of ERGIC53 and SURF4 to the ER-Golgi Intermediate Compartment and dramatic changes to the ultrastructure of the ER-Golgi interface. Disruption of TANGO1 expression also causes significant changes in early secretory pathway gene and protein expression. Disruption of both TANGO1L and TANGO1S expression impairs secretion not only of large proteins, including procollagens, but of all types of secretory cargo including small soluble proteins. Our data support a general role for Mia3/TANGO1 in maintaining both secretory pathway structure and function in vertebrate cells.

## Introduction

The first membrane trafficking step for secretion is driven by assembly of the COPII coat complex onto the endoplasmic reticulum (ER) membrane. In yeast and many other eukaryotes, this results in COPII vesicles that bud from the ER membrane (Bednarek et al., 1995). This process can be reconstituted in vitro using synthetic liposomes and a minimal COPII machinery of a small GTP binding protein Sar1p that, in its GTP-bound form, recruits an inner coat of Sec23p-Sec24p, and subsequently an outer coat of Sec13p-Sec31p (Matsuoka et al., 1998). Together these proteins are sufficient to generate 60-80 nm vesicles in an energy-dependent manner. Several other proteins support the COPII system including the guanine nucleotide exchange factor Sec12p that activates Sar1p, and Sec16p which potentiates vesicle formation (Espenshade et al., 1995). In metazoans, COPII proteins, including Sec16, assemble at relatively stable sites on the ER membrane called transitional ER (Orci et al., 1994) from which COPII vesicles bud. In the most commonly accepted models, these vesicles then coalesce to form an ER-Golgi Intermediate Compartment (ERGIC, (Schweizer et al., 1988)). Collectively these structures form ER exit sites (ERES, (Hughes et al., 2009)). These sites are the location for cargo selection and bud formation. ERES are organized by Trk-fused gene (TFG) which forms a meshwork around nascent budding sites (Johnson et al., 2015; McCaughey et al., 2016; Witte et al., 2011). Apoptosis-linked gene 2 (ALG-2) promotes oligomerization of TFG (Kanadome et al., 2017). The Sec23-interacting protein, SEC23IP (also known as p125) is a lipid binding protein that acts to promote COPII budding (Klinkenberg et al., 2014; Tani et al., 1999). Despite the prevalence of this vesicular model in the literature, few studies have identified any significant number of 60-80nm secretory vesicles in vertebrate cells or tissues. While they have been detected (Martinez-Menarguez et al., 1999) they are not abundant.

In metazoans, many proteins have been identified which help orchestrate and regulate ERES membrane dynamics in diverse ways. TANGO1, encoded by Mia3, was identified in a genetic screen in *Drosophila* S2 cells, as a factor required for the secretion of a horseradish peroxidase reporter (Bard et al., 2006). Considerable data have since implicated this as a selective cargo receptor for procollagens (Maeda et al., 2016; Raote et al., 2020; Raote et al., 2018; Saito et al., 2011; Santos et al., 2015) and other large cargo proteins such as apolipoproteins (Santos et al., 2016). Knockout of TANGO1 in *Drosophila* leads to defects in ER morphology, induction of ER stress, and defects in cargo secretion (Rios-Barrera et al., 2017). In *Drosophila*, many cargoes are TANGO1-dependent (Liu et al., 2017) including type IV collagen, the sole collagen expressed. *Drosophila* do not produce a fibrillar collagen matrix but do secrete some larger cargoes such as Dumpy (Rios-Barrera et al., 2017). Notably *C. elegans* do not express TANGO1 but TMEM131 (Zhang et al., 2020) and/or TMEM39 (Zhang et al., 2021) might play a similar role in collagen secretion in nematodes.

Analysis of TANGO1 function in vertebrates is complicated by the presence of multiple isoforms encoded by the Mia3 gene, the two principal ones being TANGO1S and TANGO1L indicating the short (785 amino acid) and long (1907 amino acid) forms respectively (McCaughey and Stephens, 2019). The longer form contains an ER luminal SH3 domain that is reported to engage cargo, including procollagen by binding to the chaperone Hsp47 (Ishikawa et al., 2016). Notably, evidence from RNAi experiments suggests that TANGO1L and TANGO1S function interchangeably in terms of procollagen secretion (Maeda et al., 2016). TANGO1 recruits Sec16 (Maeda et al., 2017) and proteins encoded by Mia2 gene including cTAGE5 (cutaneous T cell lymphoma-associated antigen 5, (Ma and Goldberg, 2016; Saito et al., 2011)). Together cTAGE5 and TANGO1 form larger complexes with Sec12 (Maeda et al., 2016), the guanine nucleotide exchange factor that promotes assembly of COPII via GTP loading of Sar1. TANGO1 also integrates with SNARE proteins to recruit membranes from the ER-Golgi intermediate compartment (ERGIC) to ERES (Nogueira et al., 2014). This has been considered to provide additional membrane to promote bud expansion to facilitate encapsulation of large cargo such as fibrillar procollagens (Ma and Goldberg, 2016; Nogueira et al., 2014; Raote et al., 2018; Saito et al., 2009) and pre-chylomicrons (Santos et al., 2016) in ‘mega-carriers’. This model has been extended following the identification of ring-like structures of TANGO1 that could be consistent with the “neck” of an emerging bud (Liu et al., 2017; Raote et al., 2017). TANGO1 is clearly a key component of the ER export machinery for large cargo (Raote and Malhotra, 2021) but questions remain as to its more general role in membrane trafficking and the contribution of the different isoforms.

A Mia3 knockout mouse (Wilson et al., 2011) has been described that has substantial defects in bone formation leading to neonatal lethality. Biallelic mutations in TANGO1 have been described in humans that result in skipping of exon 8 and multiple defects including skeletal abnormalities, diabetes, hearing loss and mental retardation (Lekszas et al., 2020). This is reflected in TANGO1 mutant zebrafish models where multiple organs were found to be affected following loss of TANGO1 or its closely related orthologue, cTAGE5 (Clark and Link, 2021). Total loss of TANGO1 expression in humans is perinatally lethal showing an absence of bone mineralization (Guillemyn et al., 2021), reflecting the phenotype seen in Mia3^-/-^ mice (Wilson et al., 2011).

Here, we have used CRISPR-Cas9 genome engineering of human cells to disrupt expression of either the short form or both short and long forms of TANGO1 to define the impact on secretory pathway organization and function. Our data show only minimal changes on loss of TANGO1L but substantial impacts on the ER-Golgi interface following near-complete reduction of both TANGO1S and TANGO1L. Although cells are still viable, this results in major changes in ultrastructure, secretory function, and notably expression of genes encoding key secretory pathway machineries. These defects correlate with the severity of gene disruption. Together our data define a core requirement for TANGO1 in secretory pathway function beyond that of its role as a receptor for procollagen and other large cargo.

## Results

### Validation of Mia3 disruption

To investigate the relative contribution of TANGO1S and TANGO1L at the ER-Golgi interface we generated knockout human cell lines using CRISPR-Cas9. We designed guide RNAs against exon 2 to knockout TANGO1L and against exon 7 to knockout both TANGO1S and TANGO1L (Fig. 1A). The outcome from genome sequencing of these clones is shown in Fig. 1B. We were not able to design gRNAs to selectively target TANGO1S owing to the shared sequence with TANGO1L. Clonal mutant cell lines were validated using antibodies selective for TANGO1L or both TANGO1S and L by immunoblotting (Fig. 1C) and immunofluorescence (Fig. 1D). We obtained clones that lacked expression of TANGO1L only (denoted TANGO1 L-/S+), clones in which the transmembrane domain encoded by exon 7 as well as TANGO1S were absent but the remaining truncated TANGO1L protein was expressed (TANGO1 LΔ/S-), and clones in which both TANGO1L and TANGO1S were absent or the expression of the truncated TANGO1L drastically reduced (TANGO1 L-/S-). We present data from two clones of each but note that these differ in subtle but important ways. The trace amounts of protein persisting in the clones could be due to lines not being completely clonal or some residual expression within some cells (Fig. 1C). TANGO1 L-/S- clone 1 does not express detectable TANGO1S and only trace levels of TANGO1L and therefore is the closest to a complete TANGO1 knockout cell line. TANGO1 L-/S- clone 2 is quite heterogenous indicating a possible lack of true clonality. TANGO1 LΔ/S- clone 2 and TANGO1 L-/S- clone 1 grew very slowly (approximately 5-fold slower than wild-type (WT)). It is also important to note that the data we present are not from different clones of the same cell line but reflect distinct edits to the MIA3 locus.

**Fig. 1.**
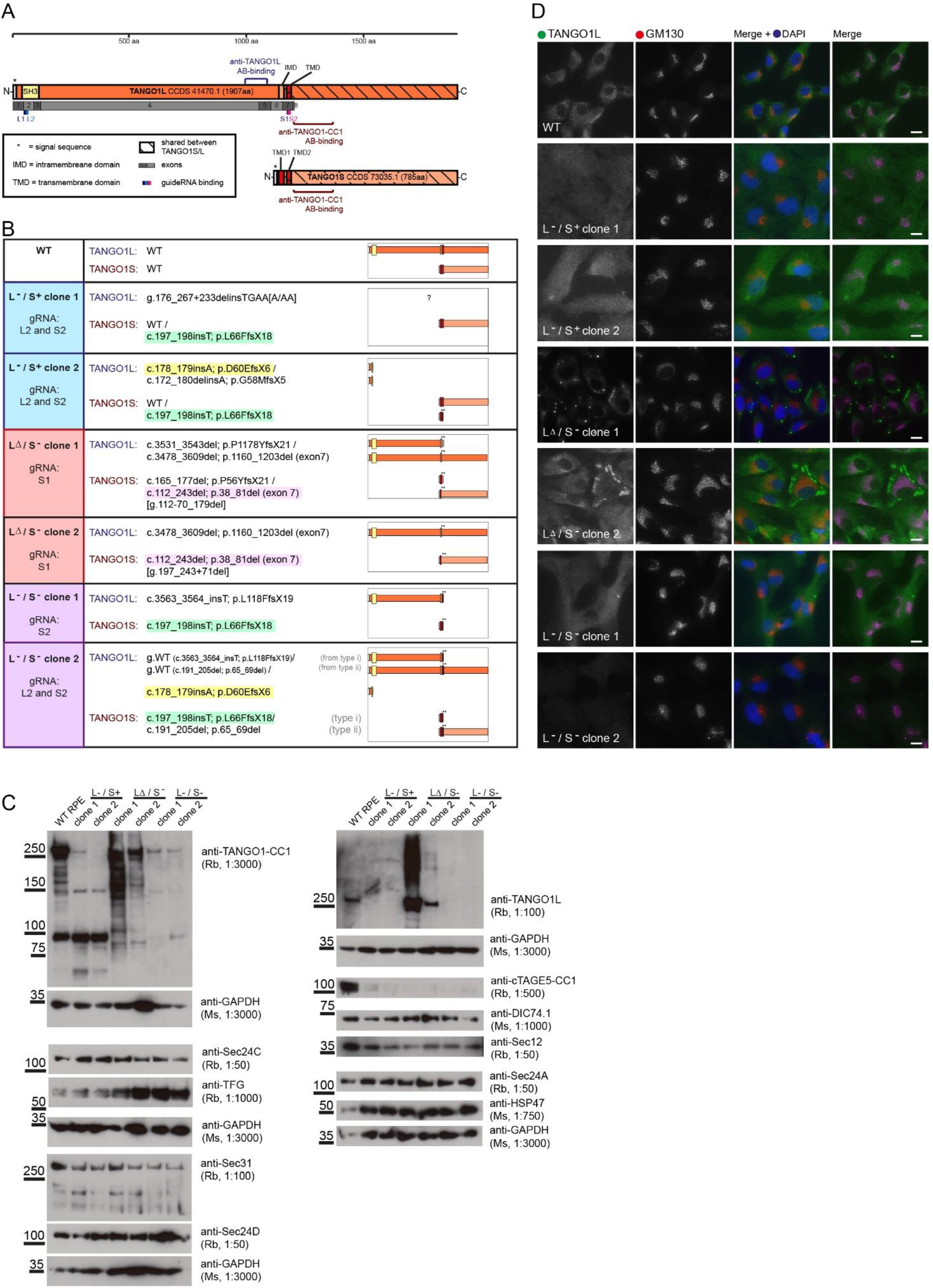
(A) Schematic showing locations of gRNAs used and antibody epitopes in relation to the major isoforms encoded by Mia3, TANGO1L and TANGO1S. (B) Mapped genomic changes including a schematic of the predicted outcomes for encoded proteins. (C) Immunoblotting using TANGO1 antibodies that detect TANGO1L and TANGO1S (anti-TANGO1-CC1) with GAPDH used as a loading control. The same lysates were used to probe for Sec24C, TFG, Sec31, Sec24D. GAPDH or DIC74.1 was used as a loading control in each case, boxes indicate the individual gels that relate to those control blots. Lysates were also probed to detect TANGO1L, cTAGE5, Sec12, Sec24A, and Hsp47. (D) Immunofluorescence was used to confirm loss of TANGO1L expression (green in merge) and co-labelled to detect the Golgi using GM130 (red in merge). Bar = 10 μm.

We used RNAseq to determine both the changes in transcription from the MIA3 locus as well as global changes in gene expression, including any possible compensation, resulting from disruption of MIA3 expression. Evident from this analysis (Fig. 2) is significant disruption to exon 1 and 2 in all clones including a near-complete loss of reads at around the start codon for TANGO1L (ENST00000344922.10) and disruption within exon 2. In clones where exon 7 was targeted, we found significant disruption within exon 7 itself and more impact on the upstream sequence (which we term exon 6a) that encodes the start site for TANGO1S (ENST00000340535.11).

**Fig. 2.**
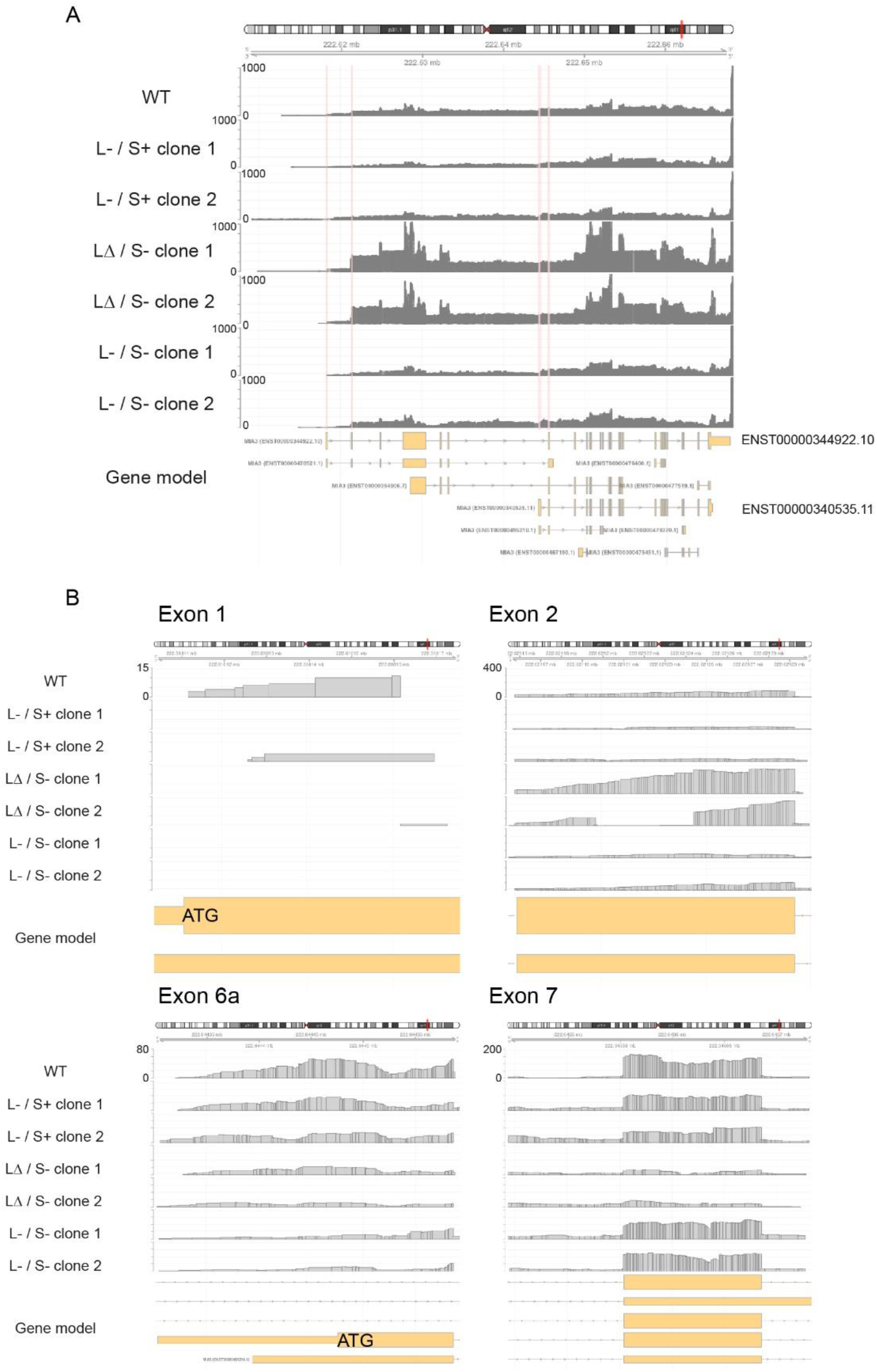
RNAseq read mapping. A. Gene model for Mia3 gene (showing chr1:222,608,000 - 222,668,400) with read depth mapped for each cell line. B. Regions of particular interest (Exon 1, chr1:222,618,104 - 222,618,176; Exon 2, chr1:222,621,155 - 222,621,301; Exon 6a, chr1:222,644,301 - 222,644,593; Exon 7, chr1:222,645,442 - 222,645,734) are displayed is a chromosome ideogram, a genomic coordinate region track, and a genome annotation track showing putative gene models for MIA3. The gene annotation depicts predicted splice forms; ENST00000344922.10 denotes the major isoform TANGO1L, ENST00000340535.11 depicts TANGO1S. B. Exon 1 is significantly disrupted in all knockout cell lines with few if any reads around the core ATG site for TANGO1L. Exon 2 is also disrupted. Exon 6A shows the ATG start codon for TANGO1S. Exon 7 is the target of other gRNAs used.

We used immunoblotting to analyze expression levels of COPII proteins and found that while Sec24A, Sec24C, Sec24D, Sec31A, and Sec12 were unaffected, TFG was upregulated in those knockout-cell lines where exon 7 of Mia3 was targeted. In all knockout cell lines, expression of cTAGE5 (encoded by Mia2) was dramatically reduced. In most cell lines, immunofluorescence (Fig. 1D) confirmed a loss of TANGO1L expression. Cell lines expressing a truncated TANGO1L (TANGO1 LΔ/S- lines) had notable abnormal, large, TANGO1-positive structures. Labelling for GM130 in all knockout cells showed obvious disruption of the Golgi consistent with fragmentation while retaining a broadly juxtanuclear location. Loss of TANGO1 also results in upregulation of key ER stress responses including expression of calnexin and IRE1α in our cells, consistent with substantial retention of ER cargoes (Supplementary Figure S1).

### Gene ontology analysis of trafficking machineries

A key interest was to determine the difference in impact of disrupting TANGO1L expression versus disrupting both TANGO1S and TANGO1L. We used gene ontology analysis of the RNAseq data sets to define those genes that were significantly upregulated in each cell line. Combining these, we find that disruption of only TANGO1L (TANGO1 L-/S+) identifies an enrichment (albeit only ~1.4-fold) for genes involved in regulation of transcription. In contrast, analysis of cells in which both TANGO1S and TANGO1L are disrupted (TANGO1 LΔ/S-), we identify enrichment of genes involved in COPII vesicle transport (9.5-fold enrichment), and trafficking to and within the Golgi apparatus (~5-fold enrichment). In the most severe cases of disruption (TANGO1 L-/S- clone 1), we identify enrichment for genes involved in intra-Golgi transport (6.5-fold), retrograde trafficking from the Golgi to ER (4.9-fold), intra-Golgi transport, COPII vesicle formation, and interestingly, zinc-ion transport (all around 5-fold enriched). The outputs from these analyses are included within Supplementary Figure S2.

We looked selectively at those genes within the RNAseq data set that are highlighted using gene ontology searches and from immunoblotting. We limited our analysis to TANGO1 L-/S- clone 1 as the most dramatically affected cell line. We set a cut-off to identify only those genes identified as changing at least 1 log_2_-fold and with a statistical significance of −log_2_ p >5. From this we found that transcription of many COPII genes was highly upregulated (Fig. 3A) including all layers of the COPII coat (Sar1A, Sar1B, Sec23A, Sec24A, Sec24D, Sec31A, TFG, and MIA2 (encoding cTAGE5)), and the known regulators of COPII function, SEC23IP (also known as p125) and ALG-2. Notably, except for TFG, these data contrast with the changes at the protein level where we see significant decreases in expression, notably of MIA2/cTAGE5 (Fig. 1C). Further to this we looked at those genes involved in retrograde trafficking and found strong upregulation of expression of KDEL receptor isoforms 2 and 3 and the lectin-type cargo receptor, ERGIC-53 encoded by LMAN1 (Fig. 3B). Analysis of genes encoding key Golgi proteins (Fig. 3C) including glycosyltransferases, trafficking complexes, and structural components of the Golgi matrix, revealed strong upregulation of ARF4 and associated GAP and GEFs, multiple tethering and fusion factors including components of the COPI coat (COPB1 and COPPB2), some components of the COG complex, multiple golgins and SNAREs. Many glycosyltransferases were also strongly upregulated, most notably GALNT5. In contrast (with one exception, ST3GAL5) multiple sialyltransferases were downregulated. In terms of a more general analysis of membrane trafficking proteins, we also found that a cohort of Rab proteins and their regulators were strongly upregulated (Fig. 3D). Together these data are consistent with activation of transcription leading to upregulation of core components of the secretory pathway. This is not however reflected in changes to the proteome. Likely a combination of transcriptional control, protein synthesis and degradation are required to maintain a functional secretory pathway following loss of MIA3 expression. Of note, these changes are only seen with any real significance when both short and long forms of TANGO1 are disrupted (especially in TANGO1L-/S- clone 1).

**Fig. 3.**
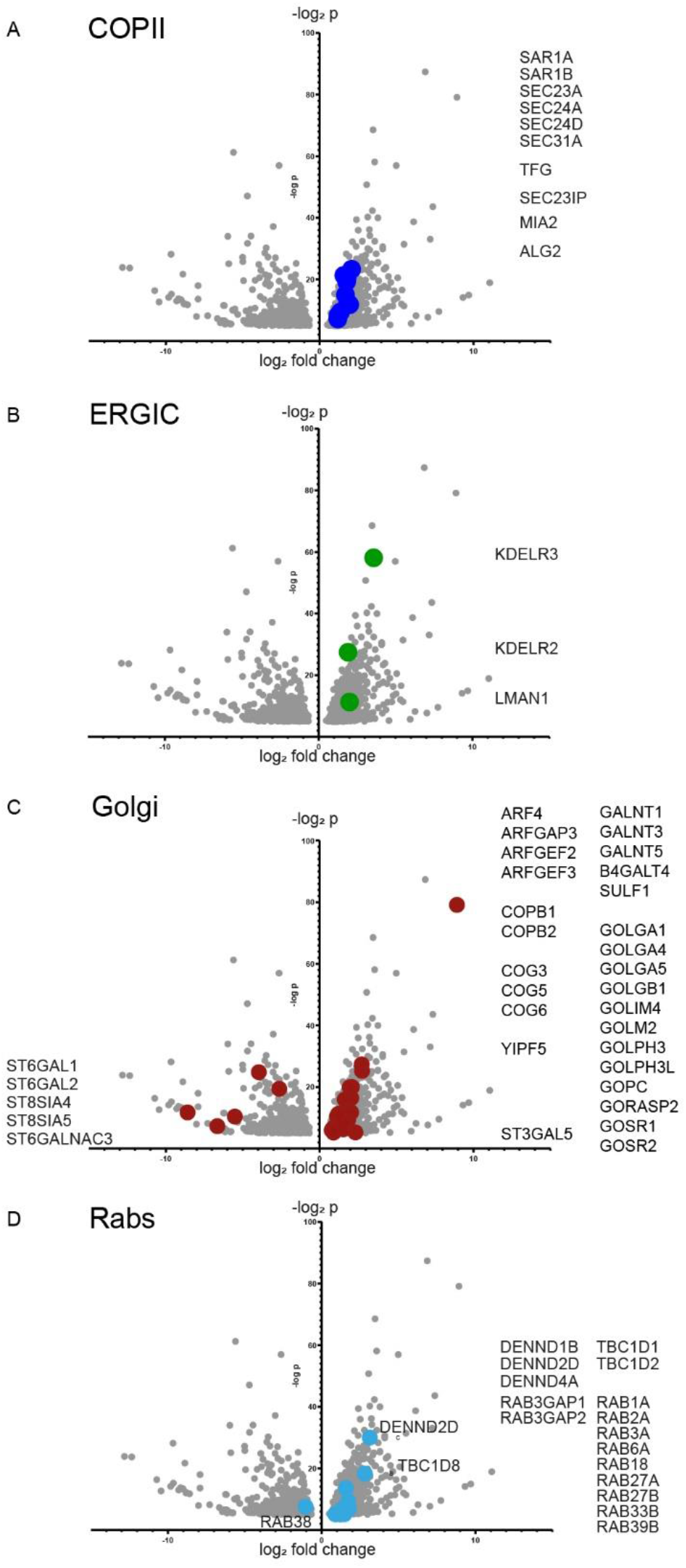
Volcano plots showing differential regulation of gene expression for (A) genes encoding COPII proteins, (B) genes encoding retrograde transport proteins, (C) genes encoding Golgi proteins, and (D) genes encoding Rabs and key regulatory proteins. Gene names listed refer to the dots highlighted on each plot. Data shows are those where -log2 p >5 and log2-fold change >1.

### Transmission electron microscopy

Given the significant changes in expression of genes encoding core secretory pathway machinery, we used transmission electron microscopy to define the ultrastructure of the ER-Golgi interface. WT cells contain a typical connected juxtanuclear Golgi ribbon (Fig. 4A). TANGO1 L-/S+ cells show disruption to Golgi structure with more separated mini-Golgi stacks evident (labelled “G” in Fig. 4B, C) with some evidence of distended ER (labelled “ER”). TANGO1 LΔ/S- cells (Fig. 4D, E) show a more severe version of this phenotype along with the presence of many small vesicular structures. TANGO1 L-/S- cells (Fig. 4F, G)) are packed with these small round vesicular structures along with other electron-lucent membranous structures that resemble degradative compartments. Scattered mini-Golgi elements are visible, as well as occasional large electron dense structures consistent with enlarged ER with retained secretory cargo (Fromme et al., 2007). The ER-Golgi interface appears very different in these cells, with fewer pleiomorphic membranes between ER and Golgi structures.

**Fig. 4.**
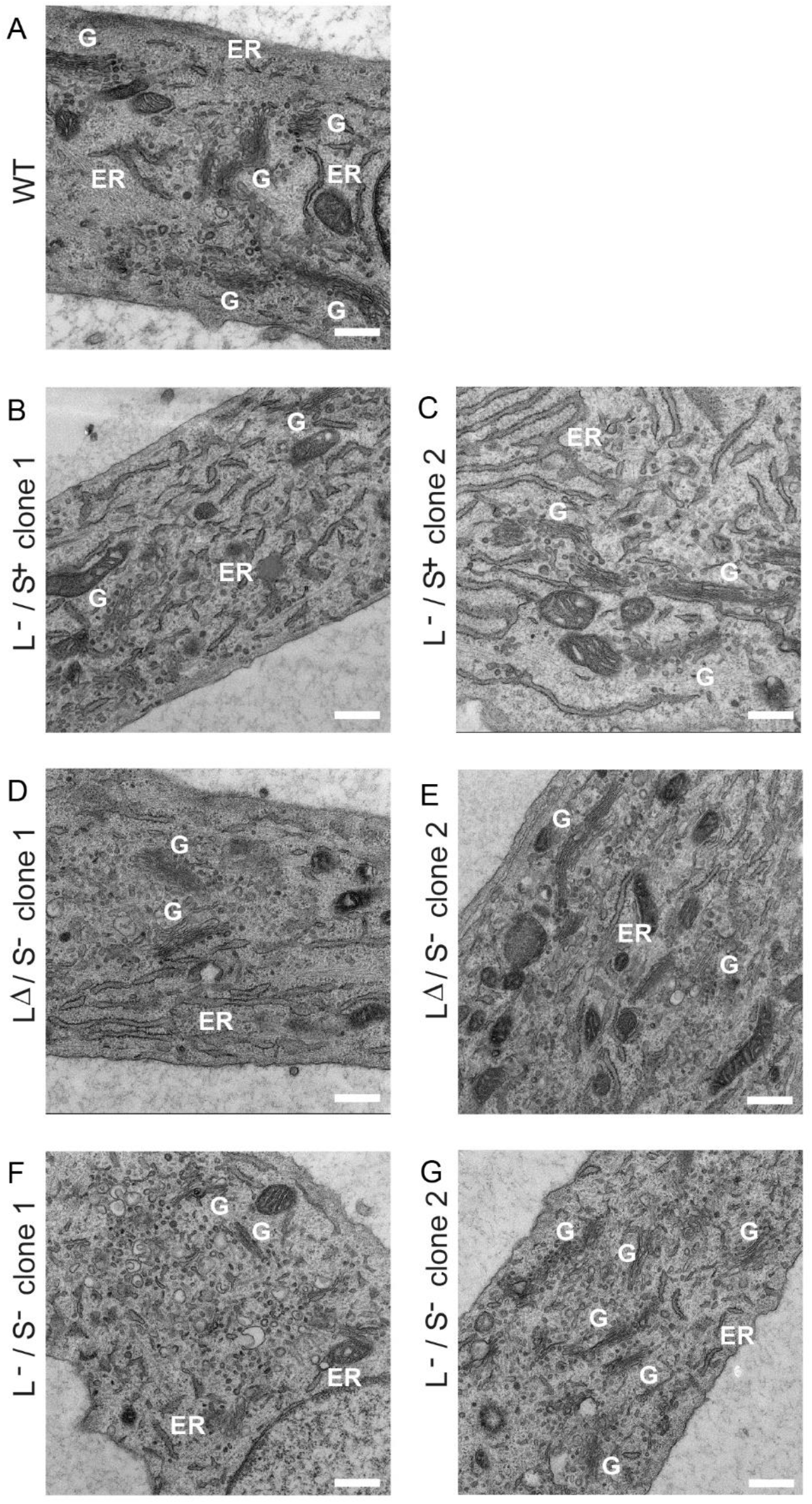
(A-G) Transmission EM was used to identify Golgi membranes in TANGO1 knockout cells compared to (A) WT. Annotations show Golgi (G) and ER membranes (ER). Bars = 500 nm.

### Phenotypic rescues by recombinant expression of TANGO1

We sought to confirm that this disruption was due to the loss of TANGO1 by reintroducing recombinant tagged proteins, TANGO1S-mScarlet-i (mSc, (Bindels et al., 2017)) or TANGO1L-HA (Raote et al., 2017)), into the knockout cells. Fig. 5 shows that TANGO1S-mSc expression reverses Golgi disruption (GRASP65 labelling, asterisks on Fig. 5A). Furthermore, TANGO1L-HA also restores a compact juxtanuclear localization of the Golgi (asterisks on Fig. 5B). Overexpression of TANGO1 isoforms often led to a more ER-like distribution than a specific ERES localization.

**Fig. 5.**
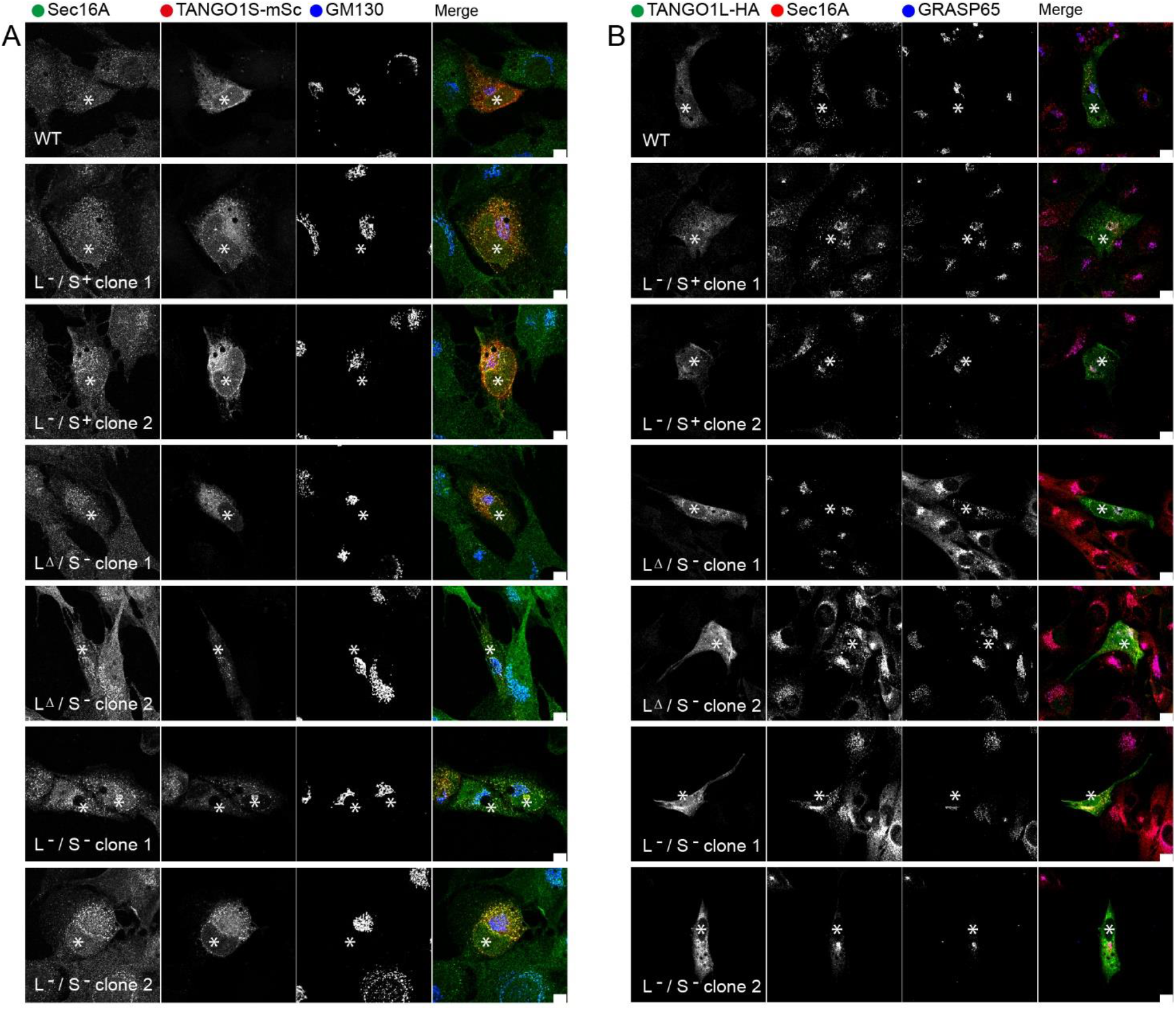
Analysis of Sec16A (A) or Sec31A (B) localization and the Golgi apparatus (GM130 (A) and GRASP65 (B)) in cells expressing TANGO1S-mScarlet-i (A) or TANGO1L-HA (B). Bar = 10 μm. Asterisks highlight cells expressing the rescue constructs.

### Disruption of ER-Golgi interface

We further analyzed the organization of the ER-Golgi interface using light microscopy (Fig. 6A and 6B, quantified in 6C-G). Fig. 6A shows localization of COPII proteins, Sec24C and Sec31A, in a characteristic punctate pattern in WT cells that is disrupted in TANGO1 knockout cells. In the most severe cases (TANGO1 L-/S-) the pattern of localization is diffuse with many more puncta detected. These changes in COPII protein distribution are also seen with Sec16A labelling (Fig. 6B). Notably COPII labelling remains clustered in a juxtanuclear position. A dramatic change in localization of ERGIC53 (Fig. 6B) is also seen with this classical marker of the ER-Golgi intermediate compartment becoming almost completely localized to the ER in TANGO1 LΔ/S- and L-/S- cells. GRASP65 and β-COP remain associated with the Golgi in all cells examined suggesting an impaired but still functional secretory pathway. Automated quantification of immunofluorescence data (Fig. 6C-G) showed an increase in the number of TFG, Sec16A, Sec24C, and Sec31A positive structures in most Mia3 knockout cell lines. This increase is consistent with the numerous vesicular structures seen by TEM, notably in TANGO1 L-/S- clone 1, and the degree of disruption correlates closely with the impact on ERGIC53 distribution. Fig. 6H shows further enlarged examples of the localization of TFG, Sec16A, and ERGIC53 in WT and L-/S- knockout cells Loss of peripheral β-COP labelling is consistent with loss of a functional ERGIC (Scales et al., 1997). We also tested the localization of another marker of the ER-Golgi intermediate compartment, surfeit 4 (SURF4, the human orthologue of the yeast cargo receptor Erv29p), in TANGO1 knockout cells. Fig. 6I shows that, like ERGIC-53, SURF4 is almost exclusively localized to the ER in TANGO1L-/S- cells. Further localization data for SURF4 in all cell lines can be found in Supplementary Figure S3.

**Fig. 6.**
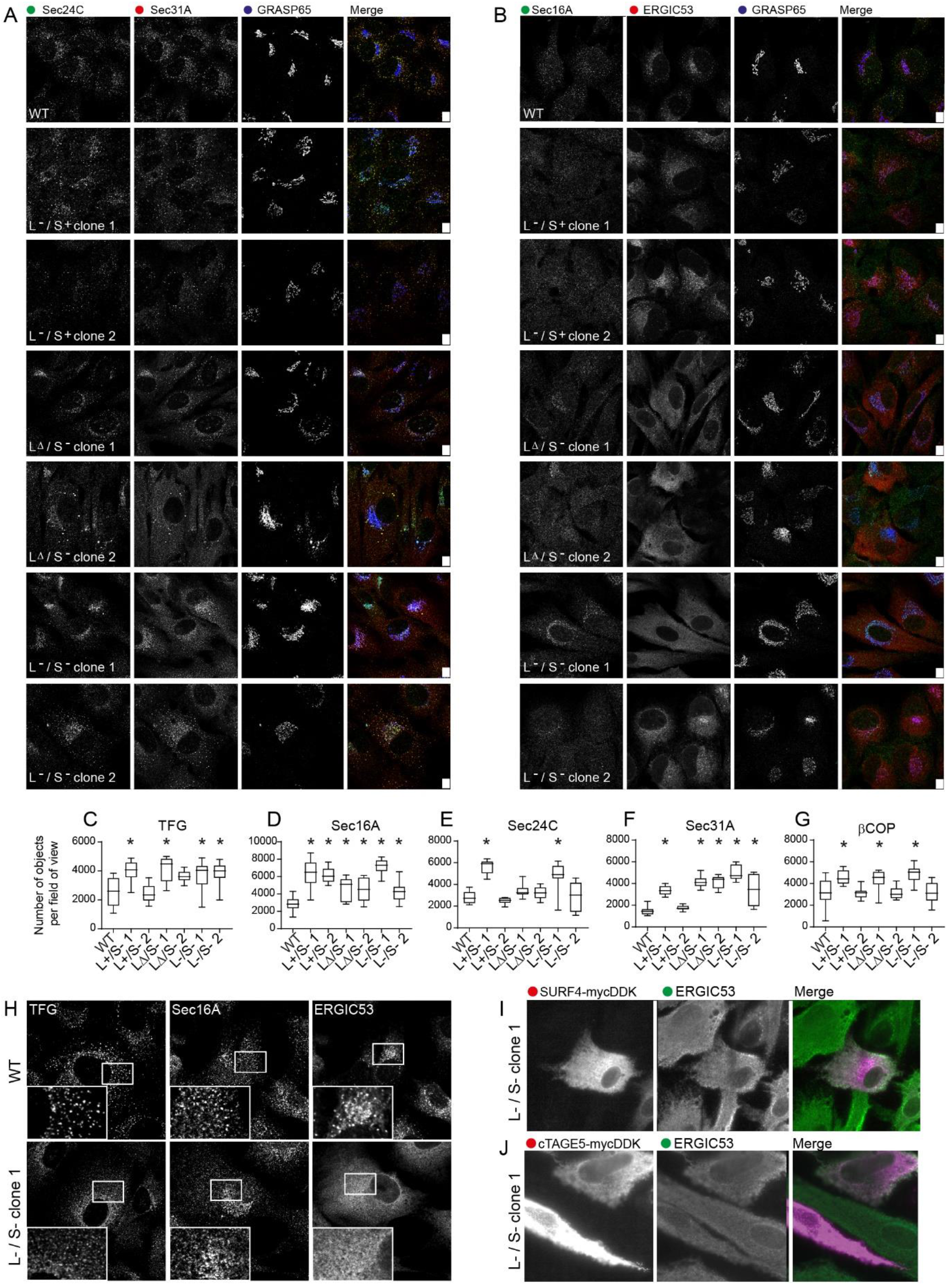
Analysis of the distribution of COPII using immunofluorescence. (A) Sec24C, Sec31A, and GRASP65 labelling. (B) Sec16A, ERGIC53, and GRASP65 labelling. Bar = 10 μm. (C-G) Quantification of these data (number of objects per field of view) for (C) TFG, (D) Sec16A, (E) Sec24C, (F) Sec31A, and (G) β-COP. Asterisks show p values <0.05 using Kruskal-Wallis tests with Dunn’s multiple comparison test for analysis of TFG and Sec24C where not all data were normally distributed, or from one-way ANOVA with Dunnett’s multiple comparison test for Sec?6A, Sec31 A and β-COP where all the data were normally distributed. (H) Enlarged views of the localization of TFG, Sed A, and ERGIC53 in WT and TANGO1 L-/S- cells. (I) Localization of SURF4-mycDDK and ERGIC53 in TANGO1 L-/S- cells. (J) Localization of cTAGE5-mycDDK and ERGIC53 in TANGO1 L-/S- cells.

Since loss of TANGO1 expression leads to concomitant loss of cTAGE5, we sought to restore the localization of ERGIC-53 in TANGO1 L-/S- cells by overexpression of cTAGE5. Overexpression of cTAGE5 did not restore the localization of ERGIC53 in TANGO1L-/S- cells to that of WT cells (Fig. 6J). The recombinant form did not exclusively localize to ER exit sites but was seen throughout the ER likely due to overexpression. Further localization data for cTAGE5 in all cell lines can be found in Supplementary Figure S4. In no cases does overexpression of cTAGE5 restore the localization of ERGIC53 to that of WT cells.

### Impact on procollagen

TANGO1 has been defined previously as a factor that selectively controls secretion of procollagens. We therefore sought to specifically test procollagen trafficking in our cell lines. Immunofluorescence shows some defects in assembly of a type I collagen matrix in our TANGO1 knockout cells, most notably in TANGO1L-/S- cells (Figure 7A). This was further supported by immunoblotting that shows intracellular retention of type I procollagen in TANGO1 knockout cells (Fig. 7B). We then used a biotin-controllable reporter to monitor procollagen transport (McCaughey et al., 2019). We were unable to derive stable cell lines of this reporter from all TANGO1 knockout clones, particularly those with the most severely disrupted Golgi morphology (LΔ/S- clone 2 and L-/S- clone 1); we interpret this as an inability of these cells to manage overexpression of procollagen in the background of an impaired secretory pathway. Unlike WT cells, all TANGO1 knockout cells were unable to transport this procollagen reporter from ER-to-Golgi within 60 minutes (Fig. 7C-G). We defined the changes in expression of all procollagen-coding genes in our cells. Fig. 8 shows that expression of many procollagens is strongly downregulated in all our Mia3-disrupted cell lines, while some show increases in expression. Of note, we detect strong downregulation of expression of COL11A1, encoding type XI procollagen, in all Mia3 knockout cell lines including where significant proportions of TANGO1L remain expressed (TANGO1LΔ/S-).

**Fig. 7.**
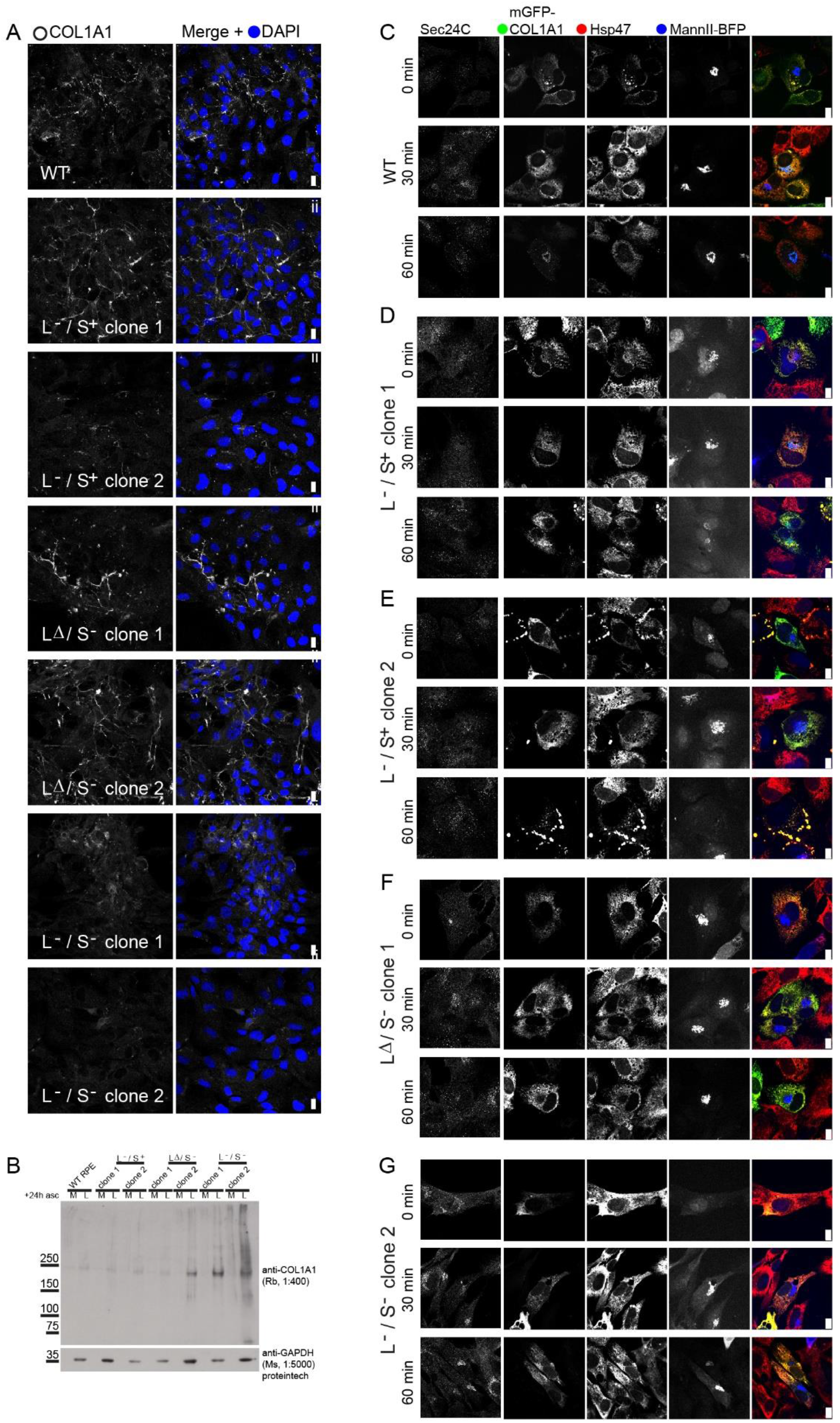
(A) Analysis of collagen I localization. Bar = 10 μm. (B) Immunoblotting to detect type I procollagen in either media (M) or lysates (L) 24h after addition of ascorbate. GAPDH is included as a loading control. (C-G) Analysis of mGFP-COL1A1 trafficking using the RUSH system. Cells were co-labeled to detect Sec24C (not included in the merge image) and Hsp47. Transfected cells co-express a Golgi marker mannll-BFP with a separate ER-hook. Bar = 10 μm.

**Fig. 8.**
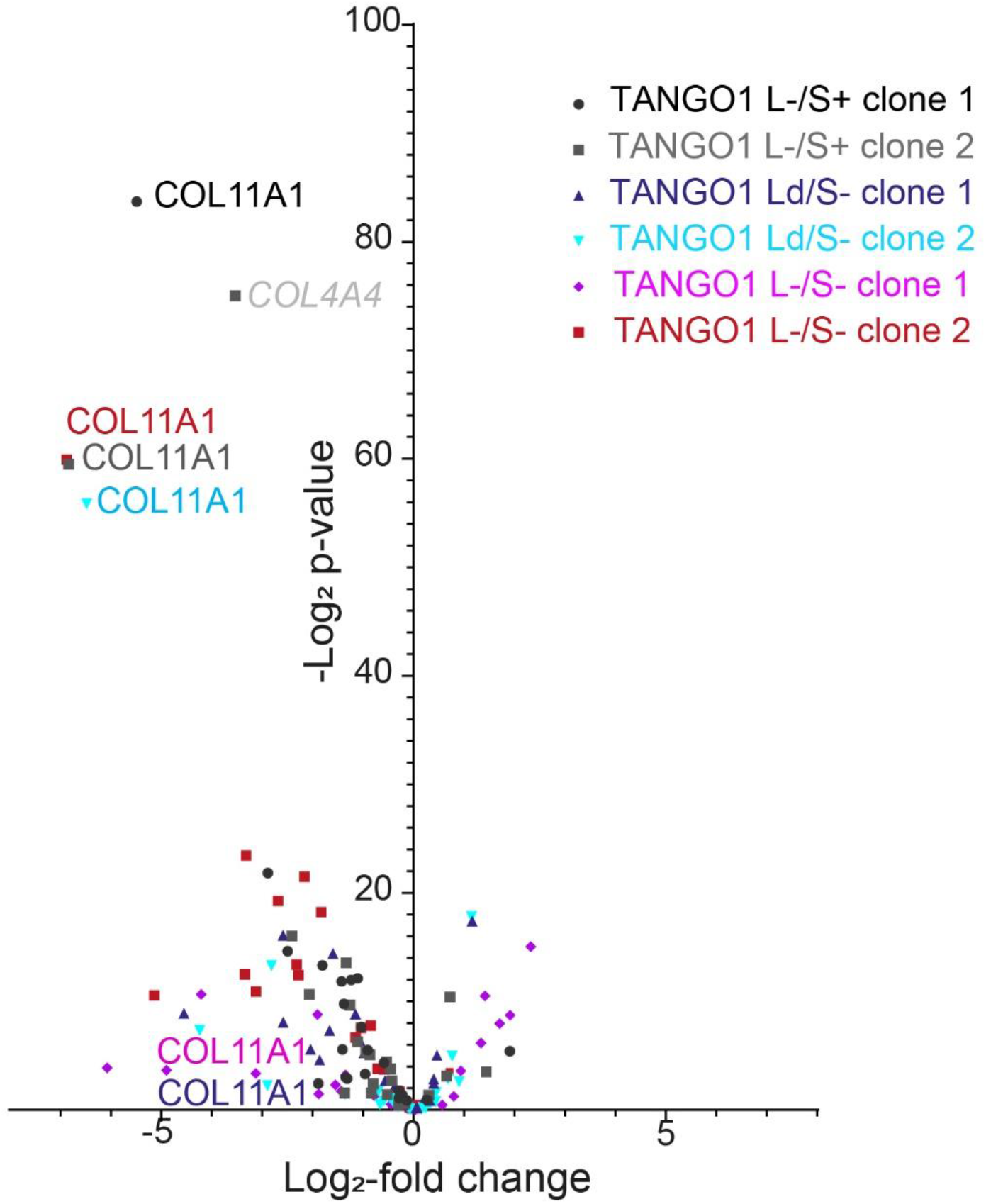
Volcano plot depicting collagen gene expression highlighting the COL11A1 gene which his most significantly disrupted. Data shows are those where -log2 p >5 and log2-fold change >1.

### Impact on general secretory trafficking

Next, we explored the efficiency of membrane traffic in both targeted and unbiased assays. Using a biotin-controlled secretion system (Boncompain et al., 2012) we monitored transfer of cargo (mannosidase II tagged with mCherry) from the ER (labelled with protein disulfide isomerase (PDI)) to the Golgi (labelled with GRASP65). At 30 minutes after addition of biotin to release mannII-mCh from the ER, around half of TANGO1 knockout cells retained mannII-mCh in the ER (Fig. 9A, B, quantified for all cell lines in C). We further explored this using E-cadherin-mCh. Here we saw a more dramatic phenotype where almost all TANGO1 LΔ/S- and L-/S- cells retained this reporter in the ER (Fig. 9D, E, quantified as green bars in Fig. 9F).

**Fig. 9.**
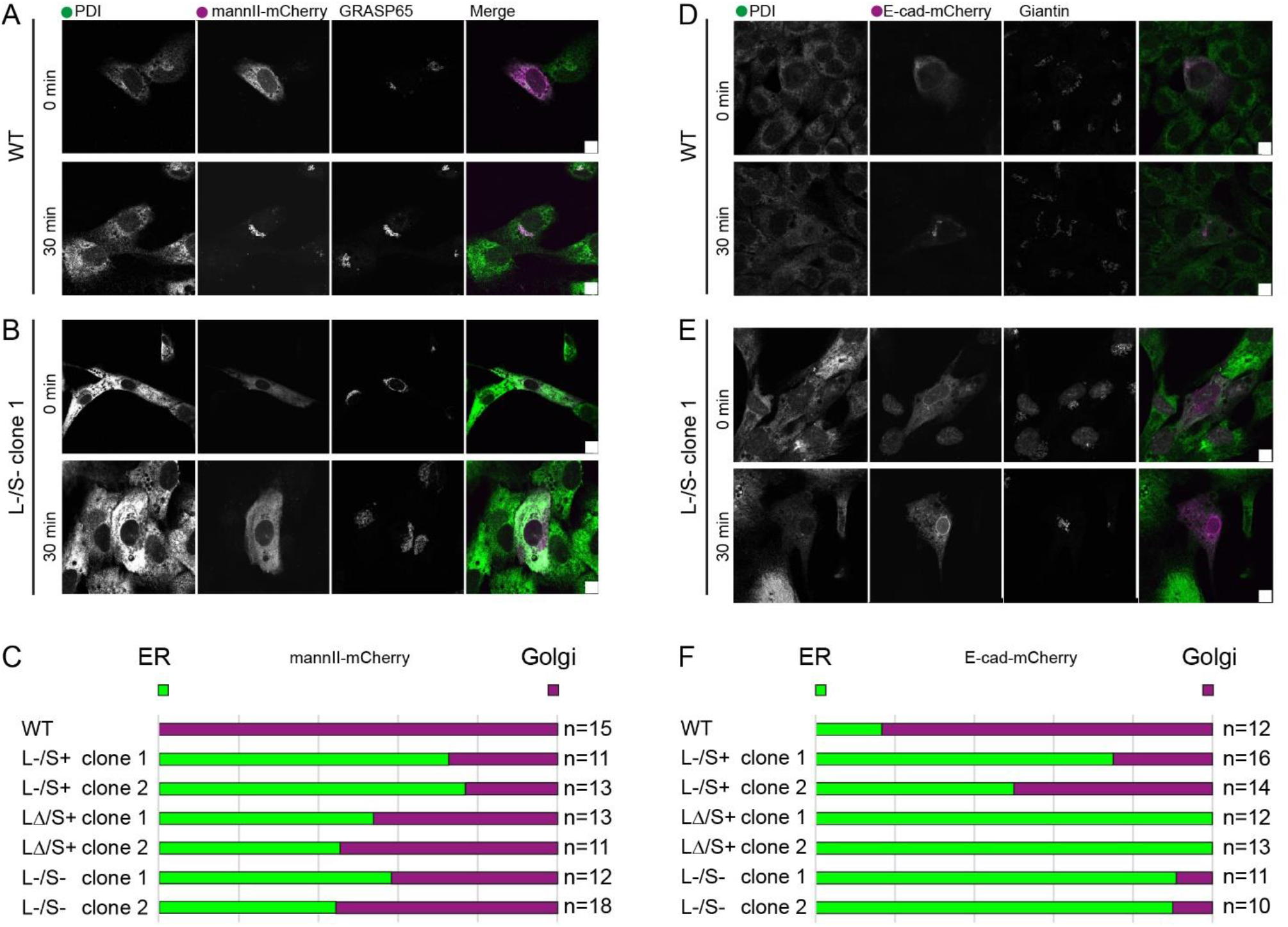
Analysis of cargo trafficking using RUSH-engineered probes (A-C) mannll-mCherry and (D-F) E-cadherin-mCherry. PDI was used to define the ER and either (A-C) GRASP65 or (D-F) giantin to define the Golgi. Bar = 10 μm. (C) These data were then analyzed to determine the number of cells in which localization of the cargo protein to either the ER (green) or Golgi (magenta) was predominant at 30 minutes after the addition of biotin.

We analyzed the secreted proteome of WT and TANGO1 knockout cells. All TANGO1 knockout cells showed defects in secretion (in all 3 biological repeats of the experiment); TANGO1 L-/S- cells showed a >5-fold reduction in secretion of amyloid precursor protein, amyloid precursor-like protein 2, NPC2, clusterin, fibrillin-1, fibrillin-2, follistatin related protein 1, IGF binding protein 7, secreted frizzled protein 1, semaphorin 7A, SERPINE, thrombospondin-1, and TGF-beta. Of note this list includes multiple components of the extracellular matrix but of varying size including many small soluble proteins. Furthermore, we saw a significant increase in secretion of fibronectin. Some of these secretion defects mirror changes at the transcriptional level including clusterin (CLU), semaphorin 7A (SEMA7A), and TGF-beta (TGFB1) which are all significantly downregulated in our RNAseq data. However, this relationship is not consistent across all cargoes when comparing the proteomic and transcriptomic data. We can conclude however that these data are not consistent with a selective defect in the secretion of large cargo.

We also analyzed the cell-derived matrix after removal of the cell monolayer by mass spectrometry. Proteins decreased at least 5-fold in abundance (and filtered for detection of >10 peptides) in TANGO1 L-/S- compared to WT cells in three independent repeats included collagen IV, protein-glutamine gamma-glutamyltransferase-2 (TGM2), fibronectin, and lysyl oxidase 1. Fibrillin-2, TGFβ, latent TGFβ binding protein 3, and perlecan were all reduced >5-fold in two of the three biological replicates. Extracellular matrix (ECM) remodeling proteins ADAMTS1 (6.9-fold), and cell migration-inducing and hyaluronan-binding protein (3.0-fold) are also decreased in all three experiments. Interestingly we also see an increase in several proteins in all three repeats including the secreted signaling proteins Wnt5b (average 7.7-fold increase), semaphorin-3C (7.2-fold) and −3D (4.0-fold), and midkine (3.8-fold). We also see increases in intracellular proteins including vimentin (5.2-fold) and interestingly, the plasma membrane integrin, ITGAV (2.6-fold). These are presumably retained on extraction of the cell layer but could importantly reflect key changes in cell architecture and adhesion. ITGAV is notable as a receptor for non-collagenous matrix ligands including fibronectin and vitronectin, notably increased in secretion to the culture medium. These data show the diversity of cargo that is affected following disruption of Mia3 expression.

## Discussion

Our data show that loss of TANGO1 expression results in dramatic morphological changes to the ER-Golgi interface coupled with significant changes in secretion. We also find that disruption of the MIA3 locus causes changes to other key machineries of the secretory pathway. Notably, we see a loss of expression of cTAGE5, a key binding partner of TANGO1 and some down-regulation of Sec31A at the protein level. We consider it likely that the reduction in cTAGE5 encoded by the Mia2 gene, is caused by interdependent stabilization of the cTAGE5-TANGO1 complex (Ma and Goldberg, 2016; Maeda et al., 2016). Notably these changes in cTAGE5-TANGO1 expression at the protein level are not reflected in our RNAseq data where in contrast we see significant upregulation of COPII and Golgi-related trafficking machines.

The specific phenotypes we define here clearly result from loss of Mia3/TANGO1 but it is difficult to ascribe a direct role owing to substantial changes in the transcriptome and proteome of edited cells. This of course could apply to any genome engineering experiment. However, key phenotypes can be rescued by restoration of either short or long forms of TANGO1 defining it as a core component of the machinery. Notably, we find no evidence that cTAGE5 can restore function to TANGO1 knockout cells. While we see a reduction in expression of COPII proteins such as Sec31A, this is not reflected at the transcriptional level where instead there is a global increase in expression of genes encoding the core secretory machinery including for both ER export and intra-Golgi trafficking. This indicates transcriptional changes as well as proteostasis mechanisms which could ensure sufficient secretory pathway activity while retaining the necessary level of quality control. This is likely linked to protein folding and stress response pathways.

A key finding here is that our data reveal only limited defects following disruption of TANGO1L alone. This includes some, but relatively minor, defects in procollagen secretion compared to those cells in which TANGO1S is also disrupted. Transcription from the Mia3 locus is strongly upregulated in cells expressing the truncated version of TANGO1L (TANGO1LΔ) likely a compensatory mechanism. The most severe defects are only seen where both TANGO1S and TANGO1L are disrupted consistent with a core role for the cytosolic domain of TANGO1 in the organization and function of the early secretory pathway. This is reflected in all assays including transcriptomic changes. Of note, the cytosolic domain of TANGO1 coordinates COPII function with that of the ERGIC (Raote et al., 2018; Santos et al., 2015). The first coiled coil of TANGO1, named TEER (Tether for ERGIC at the ER), recruits ERGIC membranes to ERES to promote membrane expansion (Santos et al., 2015). The loss of this domain in TANGO1 L-/S- cells could therefore explain the redistribution of ERGIC-53 to the ER in TANGO1S-/L- cells. Furthermore, our transcriptomic data show strong upregulation of retrograde trafficking machinery including LMAN1 (encoding ERGIC53) and two KDEL receptor isoforms KDELR2 and KDELR3. Expression of these two KDEL receptor isoforms is known to be regulated by the unfolded protein response (Trychta et al., 2018) and loss of TANGO1 in our cells does cause increases in expression of key ER stress markers. Our data also reveal strong transcriptional upregulation of the COPI trafficking machinery and particularly of ARF4, recently shown to support retrograde Golgi-to-ER trafficking (Pennauer et al., 2021). Together these data suggest an increase in retrograde trafficking as a compensatory mechanism to attenuate secretion. These data also suggest that Mia2 and Mia3 have a fundamental role in maintaining the ERGIC as a steady-state compartment, supporting models of the ERGIC as a transient organelle requiring ongoing secretory function for its formation and maintenance (Appenzeller-Herzog and Hauri, 2006).

In our engineered cell lines, notably in the most severely affected TANGO1L-/S- clone 1, we see a dramatic increase in small vesicular structures in the vicinity of ER and Golgi elements. One interpretation of these data is that TANGO1 acts normally to restrict formation of COPII vesicles, instead forming a network of ER-ERGIC contacts that include tubules to mediate more direct cargo transfer. This would be consistent with models based on rings of TANGO1 forming a neck for such structures (Liu et al., 2017; Raote et al., 2017). In addition, this could explain the paucity of COPII vesicles normally seen in mammalian cells (Martinez-Menarguez et al., 1999). TANGO1 could limit COPII vesicle formation through delaying scission of COPII coated vesicles and facilitating formation of more amorphous tubular-vesicular structures that then bud to become independent from the underlying ER. Proline-rich domains (PRDs) of TANGO1 and cTAGE5 bind to multiple copies of Sec23, initially competing with Sec31 and thereby limiting recruitment of the outer layer of the COPII coat (Ma and Goldberg, 2016). This itself would limit GTPase stimulation by Sec13-Sec31 promoting bud growth. Such a model does not require formation of “megavesicles” and could result in more tubular ER export domains. This is consistent both with previous EM analyses suggesting *en bloc* protrusion of carriers (Mironov et al., 2003) and with the lack of obvious large carriers on live cell imaging of GFP-procollagen (McCaughey et al., 2019). TFG, notably upregulated in TANGO1 knockout cells at both RNA and protein levels, could also have a key role here in generating a local environment that promotes and stabilizes this process (Johnson et al., 2015; McCaughey et al., 2016; Witte et al., 2011). This idea of direct connections as key mediators of ER-to-Golgi transport is further supported by recent high-resolution microscopy showing a network of tubules emerging from ERES to conduct secretory cargo to the Golgi (Weigel et al., 2021). Our data support models where TANGO1 promotes the maturation of membrane-bound compartments from an ER to an ERGIC and finally Golgi identity (McCaughey and Stephens, 2019).

The functions of TANGO1 in COPII coat assembly and maintenance, as well as in modulating the organization and function of the ERGIC, require the cytosolic domain of TANGO1. Additional roles for the luminal domain of TANGO1L, for example by engaging procollagen via the chaperone Hsp47 (Ishikawa et al., 2016), and binding to other ECM proteins (Ishikawa et al., 2018) might further facilitate export. We show that collagen types I, XI, and XVIII are all downregulated in Mia3 knockout cells which is consistent with a key role of Mia/TANGO1 in collagen secretion. However, the collagen isotype that is most significantly affected is type XI collagen; this suggests an alternative explanation for the gross defects in collagen secretion and collagen matrix assembly in Mia3 knockout models (Wilson et al., 2011) and patients with loss-of-function mutations in Mia3 (Guillemyn et al., 2021; Lekszas et al., 2020). Type XI collagen is a regulatory fibril-forming collagen (Birk and Brückner, 2011); the other regulatory fibril-forming collagen, type V, is not expressed at significant levels in RPE1 cells. Downregulation could be a simple and effective means to reduce the burden on cells with a compromised secretory capacity. However, these data hint at a more selective change in fibrillogenesis owing to a loss of expression of type XI collagen.

It is important to note that monitoring cell-derived matrix measures those proteins in the assembled matrix and does not necessarily reflect changes in secretion per se. Notably in both soluble proteomes and cell-derived matrix we see reductions in small soluble proteins, including TGFβ, along with many large glycoproteins of 350-500 kDa. The latter are predicted to require alternative transport mechanisms to the classical 60-80 nm COPII vesicles, although they may be sufficiently be flexible to fit (Rezaei et al., 2018) We also see increases in the presence of several key matrix proteins in cell-derived matrix that suggest cellular adaptation to defective secretory pathway function. In both proteomics data sets, changes in extracellular protein abundance do not correlate well with changes in transcription suggesting that secretory control can override increases in transcript abundance in terms of a functional matrix.

These data support models where the structural organization of the Golgi could feedback to transcriptional programmes that adapt glycosylation, possibly to ensure production of bioequivalent glycans (Mkhikian et al., 2016). Upregulation of O-glycosyltransferases including GALNT1, 3, 5 and downregulation of sialyltransferases might also result from such adaptation. Some impacts of loss of Mia3 on collagen matrix structure might reflect defects in Golgi organization that occur downstream of impacts on COPII function. However, it is notable that the increase in GALNT expression seen here contrasts with the near-complete loss of GALNT3 expression following knockout of the golgin, giantin (Stevenson et al., 2017). Loss of giantin causes only minor structural changes to the Golgi but has significant impacts on bone formation and strength (Stevenson et al., 2021). Clearly there are complex yet distinct regulatory pathways at play at both ERES and Golgi.

Together, our data show that TANGO1 plays an essential role in the organization of the ER-Golgi interface in mammalian cells and highlights the fundamental importance of endomembrane organization for effective secretion. We also show that loss of TANGO1 impacts many different types of secretory cargo in addition to procollagen. This is of course consistent with its original identification in a screen for factors affecting secretion of horseradish peroxidase (Bard et al., 2006). Overall, our data support models where ECM proteins are the cohort of secretory cargo most sensitive to perturbation of early secretory pathway function. We consider that this is likely a reflection of the extraordinary developmental secretory load during tissue development and complex glycosylation patterns of ECM proteins. Even in the absence of a collagen specific effect, the sensitivity of ECM assembly to loss of TANGO1 can still explain why bone and cartilage formation is the most dramatically affected process in Mia3 knockout *in vivo* (Wilson et al., 2011).

Our work suggests models where TANGO1 supports the formation of amorphous carriers that mediate efficient ER-to-Golgi transport of many types of protein. Our data do not support a collagen-selective role for such non-vesicular carriers in traffic to the Golgi. Instead, we suggest that the plasticity of the ER-Golgi interface underpins the efficiency of transport from the ER-to-Golgi. A diversity of routes from ER-to-Golgi including short-range and long-range intermediates could be linked to regulation of secretion for example, enabling rapid and delayed secretory responses, potentially even regulated by the circadian rhythm (TANGO1 is itself a target for circadian regulation (Chang et al., 2020)). Cell-type specific tuning of secretory capacity during differentiation could be achieved through changes in Mia3 isoform expression and/or post-translational modification to adjust both form and function of the early secretory pathway to adapt to different cargo requirements or cases of increased secretory load (Clark and Link, 2021). While much remains to be defined about the individual role of Mia2 and Mia3 gene products in cell and tissue function our work defines Mia3 as a key organizer of the ultrastructure of the ER-Golgi interface in mammalian cells and facilitator of secretory transport for diverse cargo types.

## Acknowledgements

We thank members of the Stephens lab for continued discussion and help with this work, the Wolfson Bioimaging Facility for support and advice, and Paul Martin, and Chrissy Hammond (University of Bristol, UK) and Karl Kadler, Martin Lowe, Qing-Jun Meng, Joe Swift, and Oliver Jensen (University of Manchester, UK) for helpful discussions. RNAseq was performed by the University of Bristol Genomics Facility and we thank Christy Waterfall and Jane Coghill for their help and advice.

## Funding

This work was funded by a postgraduate research scholarship from the University of Bristol to JM with additional research grant support from UKRI-MRC and UKRI-BBSRC (MR/P000177/1 and BB/T001984/1). Bioimaging equipment used in this study was funded by UKRI-BBSRC (BB/L014181/1) and through BrisSynBio, a BBSRC/EPSRC-funded Synthetic Biology Research Centre (grant number: BB/L01386X/1).

## Author contributions

JM: Investigation, Conceptualization, Data curation, Methodology, Formal Analysis, Validation, Visualization, Writing – original draft, Writing – review & editing.

NLS: Investigation (localization of SURF4, cTAGE5 rescues), Formal Analysis, Validation, Visualization, Writing – review & editing.

CRN: Investigation (EM embedding and sectioning).

JMM: Investigation (EM imaging and analysis).

AP: Formal analysis and Visualization (RNAseq data).

KH Investigation, Formal Analysis (proteomics).

DJS: Conceptualization, Data curation, Methodology, Formal Analysis, Validation, Visualization, Writing – original draft, Writing – review & editing, Funding acquisition, Project administration, Resources, Supervision, and Writing original draft: DJS.

## Competing Interests

The authors declare no competing interests. The funders had no role in the study design.

## Reagent and Data Availability

All plasmids are available through Addgene: https://www.addgene.org/David_Stephens/.

## Supplementary Figure legends

**Fig. S1.**
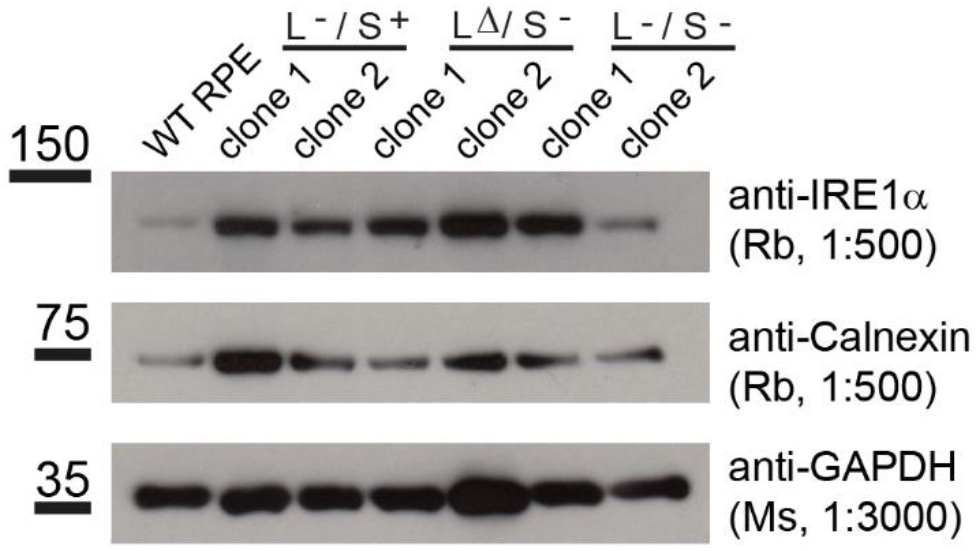
Immunoblots show expression of IRE1a, calnexin, and GAPDH (as a loading control) in TANGO1 knockout cell lines.

**Fig. S2.**
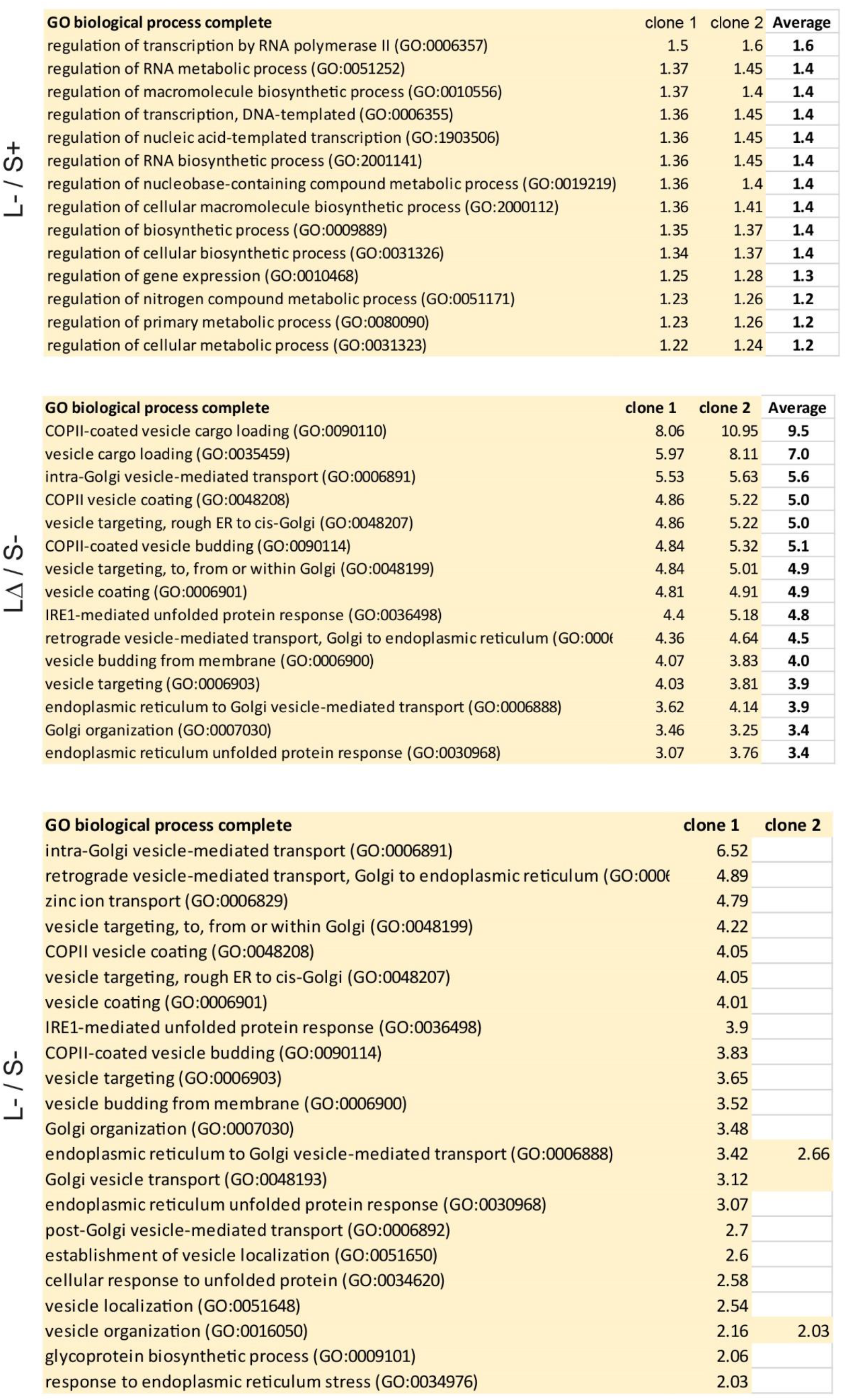
Gene ontology analysis of pooled outcomes from MIA3 knockout cell lines. Tables show those terms enriched in each cell line.

**Fig. S3.**
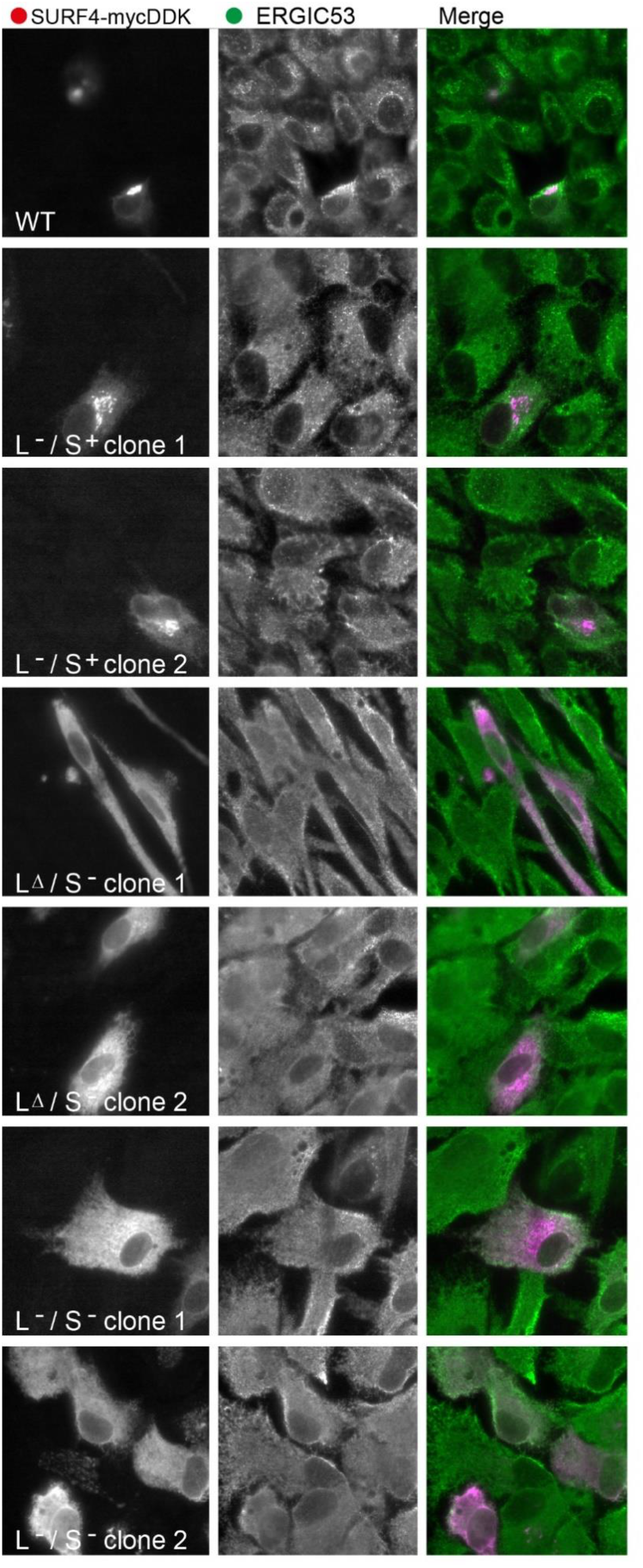
Localization of SURF4-mycDDK expressed in each cell line. Cells were transfected with SURF4-mycDDK were then fixed and processed for immunofluorescence using to detect transfected cells and endogenous ERGIC53.

**Fig. S4.**
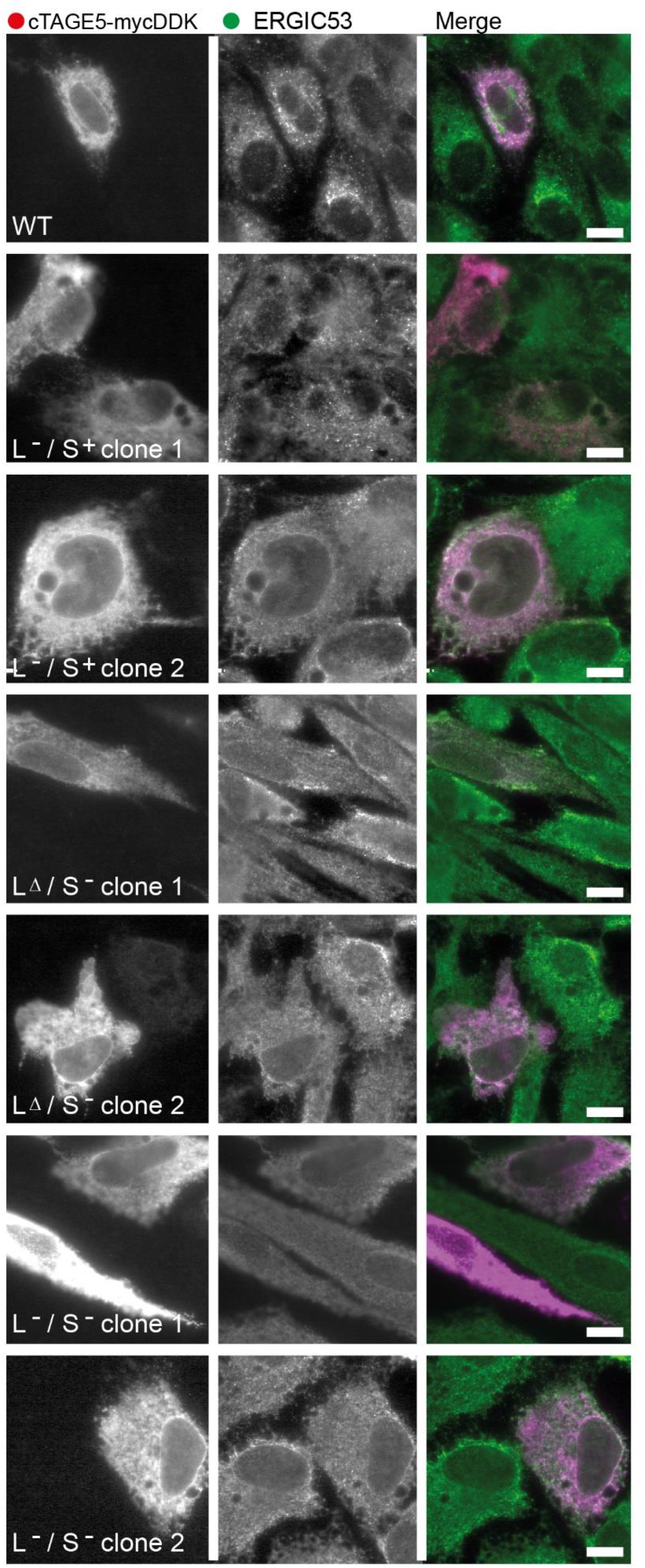
Expression of cTAGE5-mycDDK cannot restore the typical localization of ERGIC53 to TANGO1 knockout cell lines. Transfected cells were detected using the FLAG epitope tag and co-labelled to detect endogenous ERGIC53.

## Materials and Methods

All chemicals and reagents were obtained from Merck Millipore (Watford, UK) if not stated otherwise.

### Cell culture and generation of CRISPR-knockout cell lines

A human telomerase reverse transcriptase-immortalised retinal pigment epithelium type 1 cell line (hTERT RPE-1, hereafter referred to as RPE-1; ATCC^®^ CRL-4000) was used for all experiments, including the generation of stable cell lines. Cells were cultured at 37 °C and 5% CO2 in a humid environment. RPE-1 cells were grown in Dulbecco’s modified Eagle medium (DMEM) F12 supplemented with 10 % decomplemented foetal bovine serum (FBS; Thermo Fisher Scientific). Cells were passaged every 3 - 4 days when a confluence of about 80 % was reached. For passaging cells were rinsed with phosphate buffered saline (PBS) following treatment with 0.05% trypsin-EDTA (Thermo Fisher Scientific) at 37 °C until cell detachment.

The CRISPR-knockout cell lines were generated using the *TrueGuide Synthetic CRISPR gRNA* system (Thermo Fisher Scientific) with custom gRNA synthesis. GuideRNA sequences were obtained from CRISPR-knockout library against MIA3 from Sigma-Aldrich. RPE-1 cells were transfected following the manufacturer’s protocol using TrueCut™ Cas9 Protein v2 and 30 pmol gRNA duplex targeting the SH3-domain in TANGO1L (guide L1: cggtgaggctcttgaagatt or guide L2: ggattgtcgttttgtgaatt) and or the TMD in exon 7 present in both TANGO1S/L (guide S1: tgataaatacaggtttcca or guide S2: aacgaagcaattcccaaga). Cells were either transfected with only guides S1 or S2, or with both gRNAs S1+L1 or S2+L2. Clonal populations were obtained using single cell sorting and maintained in conditioned media (0.45 μm filtered one day old media supplemented with pen/strep from a confluent parent RPE-1 dish). Surviving clones were screened via immunoblotting targeting TANGO1L (2 ng per mL rabbit polyclonal anti-MIA3, Sigma-Aldrich Prestige, HPA056816-100UL) and a rabbit polyclonal targeting anti-TANGO1-CC1 (1211 – 1440aa/exon9-15 shared between both TANGO1S/L (a gift from Kota Saito, (Maeda et al., 2016))).

Stable GFP-COL1A1 expressing TANGO1 knockout cell lines were generated using virus containing the GFP-COL1A1 construct as described previously ((McCaughey et al., 2019)). In brief, the Lenti-XTM Packaging Single Shots (vesicular stomatitis glycoprotein pseudotyped version) system from Takara Bio Europe was used according to the manufacturer’s instructions (631275). Growth medium was removed from an 80% confluent 6-cm dish of RPE-1, and 1 mL harvested virus supernatant supplemented with 8 μg·mL-1 polybrene (Santa Cruz Biotechnologies) was added to cells. After 1 h of incubation at 37°C and 5% CO2, 5 mL growth medium was added. Transfection medium was then replaced with fresh growth medium after 24 h. To select for transfected cells, cells were passaged in growth medium supplemented with 15 μg·mL-1 puromycin dihydrochloride (Santa Cruz Biotechnology) 72 h after transfection and sorted via fluorescence activated cell sorting according to the signal intensity of GFP. Media for GFP-COL-RPE was further supplemented with 5 μg·mL-1 puromycin maintain engineered cell lines.

### Genotyping

Genomic DNA was obtained from clonal MIA3-knockout populations using the PureLink genomic DNA extraction kit (Thermo Fisher Scientific) according to the manufacturer’s protocol. Regions of interest were amplified by PCR using a One-Taq Hot Start DNA-polymerase (New England Biolabs) with standard buffer and 3% DMSO and the following primers targeting exon2 (L) and exon7 (S):

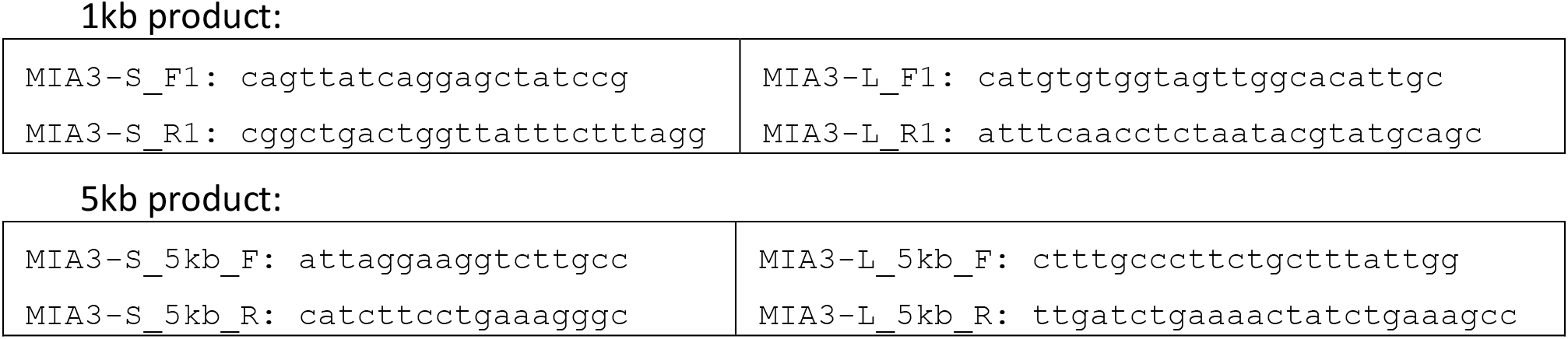
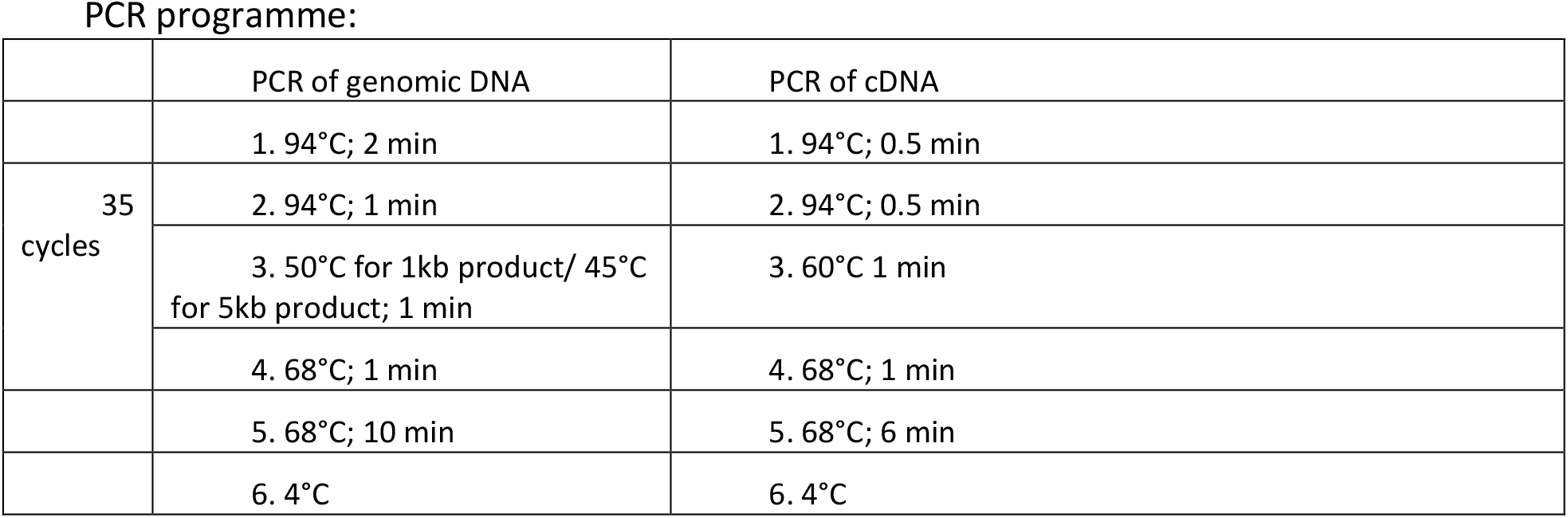

For sequence analysis of the mRNA, cDNA was generated using SuperScript III Reverse Transcription (Thermo Fisher Scientific) of extracted total mRNA using the RNeasy Plus MiniKit (Qiagen) using DTT as reducing agent. Amplification of cDNA was done as mentioned above using the following primers: MIA3-S-RT_F: ATGGCTGCGGCGCCTG, MIA3-S-RT_R: ACCTGCCACTGTGCCTTCTATCG and no supplementation of DMSO.

PCR products were ligated with the pGEM-T Easy vector system (Promega) according to the manufacturer’s protocol and transformed into DH5α *E. coli* (New England Biolabs). The plasmid DNA was extracted from 3-9 positive colonies where possible after blue-white screening using a QIAprep^®^ Spin Miniprep Kit (Qiagen) and sequenced using the standard T7 primer at Eurofins Genomics. Resulting sequences were compared using multiple sequence alignment via M-Coffee (Notredame et al., 2000) accessed at http://tcoffee.crg.cat/apps/tcoffee/do:mcoffee and displayed with the help of version 3.21 of BOXSHADE, written by K. Hofmann and M. Baron https://embnet.vital-it.ch/software/BOX_form.html.

### DNA constructs

Constructs were either generated for this study or acquired from Addgene (numbers indicated by #). The TANGO1L-HA construct was a gift from Vivek Malhotra (Saito et al., 2009). All restriction and modifying enzymes were purchased from New England Biolabs (Hitchin, UK). Str-KDEL-IRES-mannosidase II-mScarlet-i (ManII-mSc; #117274) and procollagen-SBP-mGFP-COL1A1 (#110726) were described in (McCaughey et al., 2019). StrKDEL-IRES-mannosidase II-mTagBFP2 (ManII-BFP; #165461) was generated by using ManII-mSc as a template and replacing the mScarlet-i with a mTagBFP2 generated as a synthetic gene block by Integrated DNA Technologies via restriction digest using EcoRI and FseI and subsequent HiFi NEBuilder assembly (New England Biolabs).

Synthetic gene block from IDT for generating ManII-BFP:

TCCCACCGGTCGCCACCGGaattccATGAGCGAGCTGATTAAGGAGAACATGCACATGAAGCTGTACATGGAGGGCACCGTGGACAACCATCACTTCAAGTGCACATCCGAGGGCGAAGGCAAGCCCTACGAGGGCACCCAGACCATGAGAATCAAGGTGGTCGAGGGCGGCCCTCTCCCCTTCGCCTTCGACATCCTGGCTACTAGCTTCCTCTACGGCAGCAAGACCTTCATCAACCACACCCAGGGCATCCCCGACTTCTTCAAGCAGTCCTTCCCTGAGGGCTTCACATGGGAGAGAGTCACCACATACGAAGACGGGGGCGTGCTGACCGCTACCCAGGACACCAGCCTCCAGGACGGCTGCCTCATCTACAACGTCAAGATCAGAGGGGTGAACTTCACATCCAACGGCCCTGTGATGCAGAAGAAAACACTCGGCTGGGAGGCCTTCACCGAGACGCTGTACCCCGCTGACGGCGGCCTGGAAGGCAGAAACGACATGGCCCTGAAGCTCGTGGGCGGGAGCCATCTGATCGCAAACATCAAGACCACATATAGATCCAAGAAACCCGCTAAGAACCTCAAGATGCCTGGCGTCTACTATGTGGACTACAGACTGGAAAGAATCAAGGAGGCCAACAACGAGACCTACGTCGAGCAGCACGAGGTGGCAGTGGCCAGATACTGCGACCTCCCTAGCAAACTGGGGCACAAGCTTAATggccggcctTAAggcctcgagGGCC

All other constructs for RUSH experiments were gifts from Franck Perez (Institut Curie, Paris, France: Str-KDEL-IRES-ST-SBP-mCherry (# 65265), Str-KDEL-IRES-mannosidase II-SBP-mCherry (#65253) and Str-KDEL_SBP-mCherry-Ecadherin (#65287) (Boncompain et al., 2012).

A TANGO1S-mScarlet-i (TANGO1S-mSc; Addgene #165460) construct was generated using a two-step cloning. First, StrKDEL-IRES from ManII-mSc was replaced with the human coding sequence for TANGO1S (CCDS 73035.1) via restriction digest of ManII-mSc with EcoRV and AgeI and subsequent NEBuilder HiFi assembly with the synthetic gene block containing TANGO1S with sequence overlaps at both ends. Synthetic gene block from IDT for generating TANGO1S-mSc (step 1):

aagcttggtaccgagctcggatcgatatcgcaAGCGCTgcaATGGACTCAGTACCTGCCACTGTGCCTTCTATCGCCGCTACCCCGGGGGACCCGGAACTTGTGGGACCCTTGTCTGTGCTCTACGCAGCCTTCATAGCCAAGCTGCTGGAGCTAGTTGCTACATTGCCTGATGATGTTCAGCCTGGGCCTGATTTTTATGGACTGCCATGGAAACCTGTATTTATCACTGCCTTCTTGGGAATTGCTTCGTTTGCCATTTTCTTATGGAGAACTGTCCTTGTTGTGAAGGATAGAGTATATCAAGTCACGGAACAGCAAATTTCTGAGAAGTTGAAGACTATCATGAAAGAAAATACAGAACTTGTACAAAAATTGTCAAATTATGAACAGAAGATCAAGGAATCAAAGAAACATGTTCAGGAAACCAGGAAACAAAATATGATTCTCTCTGATGAAGCAATTAAATATAAGGATAAAATCAAGACACTTGAAAAAAATCAGGAAATTCTGGATGACACAGCTAAAAATCTTCGTGTTATGCTAGAATCTGAGAGAGAACAGAATGTCAAGAATCAGGACTTGATATCAGAAAACAAGAAATCTATAGAGAAGTTAAAGGATGTTATTTCAATGAATGCCTCAGAATTTTCAGAGGTTCAGATTGCACTTAATGAAGCTAAGCTTAGTGAAGAGAAGGTGAAGTCTGAATGCCATCGGGTTCAAGAAGAAAATGCTAGGCTTAAGAAGAAAAAAGAGCAGTTGCAGCAGGAAATCGAAGACTGGAGTAAATTACATGCTGAGCTCAGTGAGCAAATCAAATCATTTGAGAAGTCTCAGAAAGATTTGGAAGTAGCTCTTACTCACAAGGATGATAATATTAATGCTTTGACTAACTGCATTACACAGTTGAATCTGTTAGAGTGTGAATCTGAATCTGAGGGTCAAAATAAAGGTGGAAATGATTCAGATGAATTAGCAAATGGAGAAGTGGGAGGTGACCGGAATGAGAAGATGAAAAATCAAATTAAGCAGATGATGGATGTCTCTCGGACACAGACTGCAATATCGGTAGTTGAAGAGGATCTAAAGCTTTTACAGCTTAAGCTAAGAGCCTCCGTGTCCACTAAATGTAACCTGGAAGACCAGGTAAAGAAATTGGAAGATGACCGCAACTCACTACAAGCTGCCAAAGCTGGACTGGAAGATGAATGCAAAACCTTGAGGCAGAAAGTGGAGATTCTGAATGAGCTCTATCAGCAGAAGGAGATGGCTTTGCAAAAGAAACTGAGTCAAGAAGAGTATGAACGGCAAGAAAGAGAGCACAGGCTGTCAGCTGCAGATGAAAAGGCAGTTTCGGCTGCAGAGGAAGTAAAAACTTACAAGCGGAGAATTGAAGAAATGGAGGATGAATTACAGAAGACAGAGCGGTCATTTAAAAACCAGATCGCTACCCATGAGAAGAAAGCTCATGAAAACTGGCTCAAAGCTCGTGCTGCAGAAAGAGCTATAGCTGAAGAGAAAAGGGAAGCTGCCAATTTGAGACACAAATTATTAGAATTAACACAAAAGATGGCAATGCTGCAAGAAGAACCTGTGATTGTAAAACCAATGCCAGGAAAACCAAATACACAAAACCCTCCACGGAGAGGTCCTCTGAGCCAGAATGGCTCTTTTGGCCCATCCCCTGTGAGTGGTGGAGAATGCTCCCCTCCATTGACAGTGGAGCCACCCGTGAGACCTCTCTCTGCTACTCTCAATCGAAGAGATATGCCTAGAAGTGAATTTGGATCAGTGGACGGGCCTCTACCTCATCCTCGATGGTCAGCTGAGGCATCTGGGAAACCCTCTCCTTCTGATCCAGGATCTGGTACAGCTACCATGATGAACAGCAGCTCAAGAGGCTCTTCCCCTACCAGGGTACTCGATGAAGGCAAGGTTAATATGGCTCCAAAAGGGCCCCCTCCTTTCCCAGGAGTCCCTCTCATGAGCACCCCCATGGGAGGCCCTGTACCACCACCCATTCGATATGGACCACCACCTCAGCTCTGCGGACCTTTTGGGCCTCGGCCACTTCCTCCACCCTTTGGCCCTGGTATGCGTCCACCACTAGGCTTAAGAGAATTTGCACCAGGCGTTCCACCAGGAAGACGGGACCTGCCTCTCCACCCTCGGGGATTTTTACCTGGACACGCACCATTTAGACCTTTAGGTTCACTTGGCCCAAGAGAGTACTTTATTCCTGGTACCCGATTACCACCCCCAACCCATGGTCCCCAGGAATACCCACCACCACCTGCTGTAAGAGACTTACTGCCGTCAGGCTCTAGAGATGAGCCTCCACCTGCCTCTCAGAGCACTAGCCAGGACTGTTCACAGGCTTTAAAACAGAGCCCATAAgcagcaACCGGTccagtgtgctggaattaattcgctgtctgcgagg

Secondly, the ManII fragment of the resulting construct was replaced with a linker region via restriction digest using EcoNI and EcoRI and NEBuilder HiFi assembly reaction to allow for direct tagging of TANGO1S with mSc.

Gene block from IDT used for the generation of TANGO1S-mSc (step2, linker):

ATGGCTCCAAAAGGGCCCCCTCCTTTCCCAGGAGTCCCTCTCATGAGCACCCCCATGGGAGGCCCTGTACCACCACCCATTCGATATGGACCACCACCTCAGCTCTGCGGACCTTTTGGGCCTCGGCCACTTCCTCCACCCTTTGGCCCTGGTATGCGTCCACCACTAGGCTTAAGAGAATTTGCACCAGGCGTTCCACCAGGAAGACGGGACCTGCCTCTCCACCCTCGGGGATTTTTACCTGGACACGCACCATTTAGACCTTTAGGTTCACTTGGCCCAAGAGAGTACTTTATTCCTGGTACCCGATTACCACCCCCAACCCATGGTCCCCAGGAATACCCACCACCACCTGCTGTAAGAGACTTACTGCCGTCAGGCTCTAGAGATGAGCCTCCACCTGCCTCTCAGAGCACTAGCCAGGACTGTTCACAGGCTTTAAAACAGAGCCCAgccgcagcagcgaattccATGGTGAGCAAGGGCGAGGC

Plasmids were amplified in DH5α E. coli (New England Biolabs) and subsequent extraction of plasmid DNA was done using a MidiPrep kit (Thermo Fisher Scientific) and 50 mL cell suspension in Lysogeny broth (LB; Thermo Fisher Scientific) with 50 μg·mL-1 ampicillin (amp). For transformation 1 ng of plasmid DNA (or 2 μL ligation reaction) was added to 50 μl thawed chemically competent 5-alpha competent E. coli (New England Biolabs) on ice, mixed gently by flicking the tube and incubated on ice for 30 min. To seal membrane openings heat shock at 42 °C was performed for 30 sec and cells were incubated on ice for 5 min. Subsequently, 450 μL super optimal broth (SOC) outgrowth medium for cell recovery was added to the transformed cells and incubated at 37 °C and 220 rounds per min (rpm) for 30 min, prior to plating on LB plates containing necessary selective antibiotics and incubated overnight at 37 °C.

For plasmid amplification, resulting colonies were picked and grown in 50 mL LB with antibiotics in suspension at 37 °C and 220 rpm overnight. Extracted plasmid DNA via PureLink kit (Thermo Fisher Scientific) performed according to the manufacturer’s instructions, with elution in 100 μL sterile filtered MilliQ H2O, was used for subsequent transfection of human cells.

For screening of bacterial colonies for presence of the correct plasmid containing the insert of interest, colonies were picked and grown in 5 mL LB with according antibiotics in suspension as mentioned above, followed by plasmid extraction via a MiniPrep kit (Qiagen) according to the manufacturer’s instructions (with an elution in 30 μL sterile filtered MilliQ H2O) and restriction digest with suitable restriction enzymes (New England Biolabs) of about 250 ng plasmid DNA for 3 h, using the corresponding protocol by New England Biolabs. DNA fragments were separated by size using gel electrophoresis of 1 – 1.5% agarose gels containing ethidium bromide running at 70 – 90 V for 40 – 50 min in Tris-acetate-EDTA (Ethylenediamine tetra acetic acid; TAE) buffer. Samples were subsequently compared on a transilluminator using UV light and positive colonies identified. Sequences were confirmed via MWG Eurofins tube sequencing services.

Mouse cDNAs encoding Surf4 (NM_011512) and cTAGE5 (NM_177321) were purchased as tagged ORF clones from Insight Biotechnology (Wembley, UK). These constructs encode C-terminal Myc-DDK tags and are cloned into pCMV6-Entry.

### RNAseq methodology and analysis

Total RNA was prepared from cells using a RNeasy^®^ Mini Kit (ThermoFisher) and assessed for integrity (RIN analysis) using the RNA Screentape assay and 2200 TapeStation system (Agilent, Stockport, UK). Each cell line was analysed in triplicate. All 21 samples had a RIN score of >8 and were taken forward to library preparation. Total RNA for each sample (100ng) was prepared into barcoded sequencing libraries using the TruSeq Stranded Total RNA kit with Ribo-Zero Plus rRNA Depletion (Illumina, Cambridge, UK) following manufacturer’s instructions. Final libraries were validated using the Agilent DNA1000 Screentape assay (Agilent, Stockport, UK) and quantified using the High Sensitivity Qubit assay (Thermofisher, UK) before equimolar normalisation and pooling. Paired end 2 x 75bp sequencing of the library pool was completed using an Illumina NextSeq500 sequencer and the Illumina High Output Version 2.5 kit, 150 cycles (Illumina, Cambridge, UK) following manufacturer’s instructions. Primary analysis was completed with Illumina RTA Software Version 2.4.11 to generate FASTQ files for analysis.

All raw reads were pre-processed for a variety of quality metrics, adaptor removal, and size selection using the FASTQC toolkit to generate high quality plots for all read libraries (http://www.bioinformatics.babraham.ac.uk/projects/fastqc). We adopted a phred30 quality cutoff (99.9% base call accuracy). RNAseq alignment and data analysis used bash and python scripting to accept RNAseq post-trimmed data as input, before ultimately producing output tables of differentially expressed transcripts. Paired-end (2×75bp) raw input data is initially trimmed for any remaining adaptors using the BBDuk suite of tools (https://jgi.doe.gov/data-and-tools/bbtools/bb-tools-user-guide/). Curated reads were then aligned with STAR to the *Homo sapiens* reference genome (GRCh38) (https://www.ncbi.nlm.nih.gov/assembly/GCF_000001405.26/). FeatureCounts is used to generate read counts, using the GRCh38 annotation for reference (Liao et al., 2014). We then used DESeq2 from the R Bioconductor package to normalise the FeatureCounts generated count matrix and call differential gene expression (DGE) via the Likelihood Ratio Test (LRT) function to compare each of the experimental groups (Love et al., 2014). Benjamini-Hochberg multiple test correction was used to produce the final P-Adjusted values that could be used for downstream data mining. Aligned reads were inspected using the Integrative Genome Viewer (IGV, (Robinson et al., 2011)). IGV enables mapping of the aligned .bam files to the GRCh38 genome and associated gene models to observe expression levels in relation to their genomic position. This enabled us to interrogate the *MIA3* gene against putative gene models to observe how expression was influenced by the CRISPR knockout. Regions of interest, as detected in IGV, were converted to BedGraph format before being plotted in the GViz R package (Hahne and Ivanek, 2016).

### Gene ontology analysis

To refine our analysis of the RNAseq data, we selected those genes with >20 reads, and selected those genes with >1 log2-fold change for analysis. We searched these lists for each cell line using geneontology.org (Ashburner et al., 2000; Gene Ontology, 2021) using the PANTHER Overrepresentation Test ((Mi et al., 2019) Released 20210224, GO Ontology database DOI: 10.5281/zenodo.4495804 Released 2021-02-01) using all *Homo sapiens* genes in the database as a reference set, with Fisher’s Exact test corrected for a false discovery rate probability of <0.05.

### Cell transfection and RUSH experiments

Cells were seeded 1 – 2 days before transfection to ensure a confluence of about 60 – 80 %. A Lipofectamine2000 transfection solution was prepared according to the manufacturer’s instructions containing a total of 0.8 (for rescue experiments with either TANGO1S-mSc or TANGO1L-HA) or 1 μg plasmid DNA per construct and 2.5 μL lipofectamine2000 (Thermo Fisher Scientific) in 200 μL OptiMEM (Thermo Fisher Scientific) per 35 mm well and was added drop wise onto the cells covered with 1 mL of fresh media. Transfected cells were incubated at culturing conditions for about 16 h prior to media change, followed by either fixation for immunofluorescence or time courses in presence or absence of culture medium supplemented with 40 μM biotin. For collagen trafficking experiments of GFP-COL1A1 cells, these were incubated first in presence of 50 μg·mL^-1^ ascorbate for 24 h and then in absence of ascorbate for 24 h prior to the trafficking experiment during which 500 μg·mL^-1^ ascorbate and 400 μM biotin was used to trigger transport from the ER to the Golgi. *Immunofluorescence*

For immunofluorescence cells were grown on 13 mm (thickness 1.5; VWR) autoclaved cover slips. Cells were washed with PBS and fixed with 4% paraformaldehyde for 15 min at RT following repeated rinsing with PBS. PFA-fixed cells were permeabilised with 0.1% (v/v) Triton-X100 for 10 min at RT and blocked with 3% bovine serum albumin (BSA) in PBS for 30-60 min. Immunolabelling with primary and secondary antibodies was performed at RT for 1 h in a humid environment and in the dark. Antibodies were diluted to the final working concentrations or dilutions as in blocking solution as follows: mouse monoclonal anti-β-COP (dilution 1:500, G61610), rabbit polyclonal anti-β’-COP (dilution 1:20, (Palmer et al., 2009)), 0.5 μg.ml^-1^ rabbit polyclonal anti-COL1A1 (NB600-408, Novus Biologicals), mouse monoclonal anti-ERGIC-53 (dilution 1:1000, clone G1/93, Alexis Biochemicals), rabbit polyclonal anti-giantin (dilution 1:2000, Poly19243, BioLegend), 0.25 μg.ml^-1^ mouse monoclonal anti-GM130 (610823, BD Bioscience), sheep polyclonal anti-GRASP65 (dilution 1:1500, a gift from Jon Lane, University of Bristol (Cheng et al., 2010)), rabbit polyclonal anti-HA-Tag (C29F4) (dilution 1:500, mAb #3724, Cell Signaling), 0.75 μg.ml^-1^ mouse monoclonal anti-Hsp47 (M16.10A1, ENZO), 0.005 μg.ml^-1^ mouse monoclonal anti-PDI (clone 2E6A11, 66422-1-Ig, Proteintech), rabbit polyclonal anti-Sec16A (dilution 1:500, KIAA0310, Bethyl Labs, Montgomery, TX), rabbit polyclonal anti-Sec24C (dilution 1:250 (Townley et al., 2008)), 0.25 μg.ml^-1^ mouse monoclonal anti-Sec31A (612350, BD Bioscience), 0.4 μg.ml^-1^ rabbit polyclonal anti-TANGO1L (HPA056816-100UL, Sigma Aldrich Prestige, 0.5 μg.ml^-1^ rabbit polyclonal anti-TFG (NBP2-24485, Novus Biologicals).

Samples were rinsed three times with PBS for 5 min after incubation with primary and secondary antibodies, respectively. As secondary antibodies 2.5 μg·mL^-1^ donkey anti-rabbit Alexa-Fluor-568-conjugated, donkey anti-mouse Alexa-Fluor-647-conjugated, or donkey anti-sheep Alexa-Fluor-488-conjugated antibodies were used (Thermo Fisher Scientific).

Samples were washed with deionised water and mounted using ProLong Diamond Antifade (Thermo Fisher Scientific) with 4’,6-diamidino-2-phenylindole (DAPI) for confocal imaging or without DAPI when transfected with ManII-BFP. For imaging via widefield microscopy, MOWIOL 4-88 (Calbiochem, Merck-Millipore, UK) mounting media was used in combination with 1 μg·mL^-1^ DAPI in PBS (Thermo Fisher Scientific) for 3 min at RT prior to repeated washing and mounting.

### Image acquisition

Images of cells transiently expressing GFP-COL and showing TANGO1L and GM130 were obtained through widefield microscopy using an Olympus IX-71 inverted microscope (Olympus, Southend, UK) as described previously (McCaughey et al., 2019).

All other immunofluorescence images were obtained with confocal microscopy using a Leica SP5II for or Leica SP8 system (TANGO1L-HA rescues only with SP8, Leica Microsystems, Milton Keynes, UK) as previously described (Saito et al., 2009). In brief, images were acquired at 400 Hz scan speed with bidirectional scanning set up, zoom factor three, three times frame averaging and, when necessary, line accumulation set to two. Fluorophores were excited with an argon laser (SP5II) or white light laser (SP8) at the required wavelengths (405, 488, 561 and 630 nm). Pixel size was chosen according to Nyquist sampling.

For immunofluorescence of extracellular COL1A1 cells were seeded near confluent on cover slips, grown for 3 days in total with the last 48 h in media with 50 μg.ml^-1^ ascorbate prior to fixation with PFA and followed by subsequent steps as described before, without permeabilization using Triton-X100. Images for extracellular collagen were acquired using a confocal SP5II system at zoom factor 1, with 3000 px, 400Hz speed, 3 times frame averaging in form of z-stacks containing three slices to capture the entire signal for collagen per field of view. Four fields of view per sample were chosen by viewing the DAPI channel only.

### Immunoblots

For semiquantitative analysis of protein levels of the COPII machinery and MIA proteins in WT-RPE-1 and MIA3-knockout clones, cells were seeded in 6-well plates and grown for 2-4 days until confluent. Cells were rinsed with ice cold PBS and lysed in 200 μL buffer containing 50 mM Tris-HCl, 150 mM NaCl, 1% (vol/vol) Triton X-100, and 1% (vol/vol) protease inhibitor cocktail (Calbiochem) at pH 7.4 on ice for 15 min and scraped using rubber policemen. Lysates were centrifuged at 13,500 rpm at 4°C for 10 min and the supernatant was denatured using LDS loading buffer and reducing agent (Thermo Fisher Scientific) at 95°C for 10 min and run under reducing conditions on a 3-8% Tris-Acetate precast gel for 135 min at 100 V in Tris-Acetate running buffer supplemented with antioxidant (Thermo Fisher Scientific). Transfer of protein bands onto a nitrocellulose membrane (GE Healthcare, Amersham, UK) was performed at 15 V overnight/ 300mA for 3-5 h or semi-dry at 90V for 1.5 h. The membrane was blocked using 5% (wt/vol) milk powder in tris buffered saline with tween20 (0.01% (vol/vol)) (TBST) for 30 min at RT and incubated with primary antibodies for 1.5 h at RT or overnight at 4°C. Primary antibody concentrations (where known) used for Wester-Blot (WB) analysis were as follows: 2.5 μg.ml-1 rabbit polyclonal anti-COL1A1 (NB600-408, Novus Biologicals), rabbit polyclonal anti-cTAGE5-CC1 (a gift from Kota Saito (Maeda et al., 2016)), mouse monoclonal anti-DIC74.1 (MAB1618, Merck), 0.33 μg.ml-1 mouse monoclonal anti-GAPDH (AM4300, Thermo Fisher Scientific) or 0.2 μg.ml-1 mouse monoclonal anti-GADPH clone 1E6D9 (60004-1-Ig, Proteintech), 0.75 μg.ml-1 mouse monoclonal anti-Hsp47 (M16.10A1, ENZO), rabbit polyclonal anti-Sec12 (a gift from the Balch Lab, (Weissman et al., 2001)), rabbit polyclonal anti-Sec24A (Satchwell et al., 2013), rabbit polyclonal anti-Sec24C (Townley et al., 2008), rabbit polyclonal anti-Sec24D (Palmer et al., 2005)), rabbit polyclonal anti-Sec31A (Townley et al., 2008), 2 μg.ml-1 rabbit polyclonal anti-TANGO1L (HPA056816-100UL, Sigma Aldrich Prestige), rabbit polyclonal anti-TANGO1-CC1 (a gift from Kota Saito, (Maeda et al., 2016)), 0.5 μg.ml-1 rabbit polyclonal anti-TFG (NBP2-24485, Novus Biologicals).

After repeated rinsing with TBST, the membrane was incubated for 1.5 h at RT with HRP-conjugated antibodies diluted in the blocking solution (1:5,000) against mouse (Jackson ImmunoResearch, AB_2340770) and rabbit (Jackson ImmunoResearch, AB_10015282), respectively. The wash step was repeated, and detection was performed using Promega enhanced chemiluminescence reaction reagents and autoradiography films (Hyperfilm MP, GE Healthcare) with 3 sec – 30 min exposure and subsequent development.

For analysis of type I collagen secretion cells were incubated in 1 mL serum-free culture medium supplemented with or without 50 μg·mL^-1^ ascorbate for 24 h and the media fractions collected prior to cell lysis as described above without scraping to obtain lysis fractions. The transfer onto the nitrocellulose membrane was performed overnight at 15 V.

### Electron microscopy

WT and MIA3-knockout cell lines were grown until confluent, prior to rinsing with serum-free culture medium and subsequent fixation in 2.5% glutaraldehyde in serum free medium for 30 mins at room temperature. Cells were osmicated using osmium ferrocyanide following standard procedures and removed from the culture surface with a rubber policeman. Cells were mixed and spun down into a BSA gel at room temperature. The encased cells were then dehydrated in an alcohol series and embedded in EPON, sections (70 nm) were stained with uranyl acetate and lead citrate and imaged in a Thermo Fisher Tecnai 12 BioTwin TEM operating at 120kV with images recorded on a Thermo Fisher CETA 4kx4k camera. At least 15 cells from each variant were imaged with every sectioned Golgi complex in each cell being imaged. To aid accurate quantification, section tilting was used to enable imaging of the Golgi membranes as close to perpendicular as possible in the electron beam.

### Sample preparation and analysis of proteomes via mass spectrometry

Soluble secreted proteomes were obtained from a 6-well dish with a confluent layer of cells incubated in presence of 50 μg·mL-1 ascorbate for 24 h in 1 ml serum free media prior to sample collection. Media fractions were collected, and potential cell debris removed by centrifugation at 13500 rpm for 10 min at 4C. Samples were frozen prior to further processing for tandem-mass-tagging (TMT).

To obtain the proteome from the cell-derived matrix, cells were grown in 15 cm dishes for seven days in media supplemented with 50 μg·mL-1 ascorbate (media was refreshed every 72 hours). All clones were seeded to reach confluency on day 3. Upon sample collection cells were rinsed with ice-cold PBS and extracted in 8 mL 20 mM ammonium hydroxide and 0.5% triton-X-100 in PBS for 2 min on a shaker with subsequent repeated rinsing in ice-cold PBS and incubation with 10 μg·mL-1 DNAseI for 30 min at 37C. Samples were washed with deionised water twice and directly scraped using a rubber-policeman in 400 μL reducing agent containing LDS buffer (NuPAGE, Thermo Fisher Scientific) and boiled at 95C for 10 min prior to snap freezing in liquid nitrogen and storage at −80C until further processing for TMT.

### TMT Labelling and High pH reversed-phase chromatography

For the secretome analysis, each media sample was concentrated to approximately 100ul using a centrifugal filter unit with a 3kDa cut-off (Merck Millipore, Cork, Ireland), digested with trypsin (2.5μg trypsin; 37°C, overnight) and labelled with Tandem Mass Tag (TMT) ten plex reagents according to the manufacturer’s protocol (Thermo Fisher Scientific, Loughborough, UK), and the labelled samples pooled.

For the cell-derived matrix analysis, each sample was loaded onto a 10% SDS-PAGE gel and electrophoresis performed until the dye front had moved approximately 1 cm into the separating gel. Each gel lane was then cut into a single slice and each slice subjected to in-gel tryptic digestion using a DigestPro automated digestion unit (Intavis Ltd.). The resulting peptides were quantified using a quantitative colorimetric peptide assay kit (Pierce/Thermo Scientific) and an equal amount of each labelled with Tandem Mass Tag (TMT) ten plex reagents according to the manufacturer’s protocol (Thermo Fisher Scientific) and the labelled samples pooled.

For both secretome and cell-derived matrix analyses, the TMT-labelled pooled samples were desalted using a SepPak cartridge according to the manufacturer’s instructions (Waters, Milford, Massachusetts, USA). Eluate from the SepPak cartridge was evaporated to dryness and resuspended in buffer A (20 mM ammonium hydroxide, pH 10) prior to fractionation by high pH reversed-phase chromatography using an Ultimate 3000 liquid chromatography system (Thermo Fisher Scientific). In brief, the sample was loaded onto an XBridge BEH C18 Column (130Å, 3.5 μm, 2.1 mm X 150 mm, Waters, UK) in buffer A and peptides eluted with an increasing gradient of buffer B (20 mM Ammonium Hydroxide in acetonitrile, pH 10) from 0-95% over 60 minutes. The resulting fractions were concatenated to generate a total of four fractions, which were evaporated to dryness and resuspended in 1% formic acid prior to analysis by nano-LC MSMS using an Orbitrap Fusion Lumos mass spectrometer (Thermo Scientific).

### Nano-LC Mass Spectrometry

High pH RP fractions were further fractionated using an Ultimate 3000 nano-LC system in line with an Orbitrap Fusion Lumos mass spectrometer (Thermo Scientific). In brief, peptides in 1% (vol/vol) formic acid were injected onto an Acclaim PepMap C18 nano-trap column (Thermo Scientific). After washing with 0.5% (vol/vol) acetonitrile 0.1% (vol/vol) formic acid peptides were resolved on a 250 mm × 75 μm Acclaim PepMap C18 reverse phase analytical column (Thermo Scientific) over a 150 min organic gradient, using 7 gradient segments (1-6% solvent B over 1min., 6-15% B over 58min., 15-32%B over 58min., 32-40%B over 5min., 40-90%B over 1min., held at 90%B for 6min and then reduced to 1%B over 1min.) with a flow rate of 300 nl min-1. Solvent A was 0.1% formic acid and Solvent B was aqueous 80% acetonitrile in 0.1% formic acid. Peptides were ionized by nano-electrospray ionization at 2.0kV using a stainless-steel emitter with an internal diameter of 30 μm (Thermo Scientific) and a capillary temperature of 300°C.

All spectra were acquired using an Orbitrap Fusion Lumos mass spectrometer controlled by Xcalibur 3.0 software (Thermo Scientific) and operated in data-dependent acquisition mode using an SPS-MS3 workflow. FTMS1 spectra were collected at a resolution of 120 000, with an automatic gain control (AGC) target of 200 000 and a max injection time of 50ms. Precursors were filtered with an intensity threshold of 5000, according to charge state (to include charge states 2-7) and with monoisotopic peak determination set to Peptide. Previously interrogated precursors were excluded using a dynamic window (60s +/-10ppm). The MS2 precursors were isolated with a quadrupole isolation window of 0.7m/z. ITMS2 spectra were collected with an AGC target of 10 000, max injection time of 70ms and CID collision energy of 35%.

For FTMS3 analysis, the Orbitrap was operated at 50 000 resolution with an AGC target of 50 000 and a max injection time of 105ms. Precursors were fragmented by high energy collision dissociation (HCD) at a normalized collision energy of 60% to ensure maximal TMT reporter ion yield. Synchronous Precursor Selection (SPS) was enabled to include up to 10 MS2 fragment ions in the FTMS3 scan.

### Proteomic Data Analysis

The raw data files were processed and quantified using Proteome Discoverer software v2.1 (Thermo Scientific) and searched against the UniProt Human database (downloaded August 2020: 167789 entries) using the SEQUEST algorithm. Peptide precursor mass tolerance was set at 10ppm, and MS/MS tolerance was set at 0.6Da. Search criteria included oxidation of methionine (+15.995Da), acetylation of the protein N-terminus (+42.011Da) and Methionine loss plus acetylation of the protein N-terminus (−89.03Da) as variable modifications and carbamidomethylation of cysteine (+57.021Da) and the addition of the TMT mass tag (+229.163Da) to peptide N-termini and lysine as fixed modifications. Searches were performed with full tryptic digestion and a maximum of 2 missed cleavages were allowed. The reverse database search option was enabled and all data was filtered to satisfy false discovery rate (FDR) of 5%.

### Data analysis and statistics

Identification and analysis of objects in immunofluorescence images in Figure 4 was conducted in Volocity version 6.3 followed by statistical analysis using GraphPad Prism version 8. Data distribution was assessed for normality and then analysed using the Kruskal-Wallis test with Dunn’s multiple comparison for non-parametric data or ordinary one-way ANOVA with Dunnett’s multiple comparisons test for parametric data. Asterisks on plots indicate p-values <0.05. RUSH assays were scored by visual inspection of predominant localization of reporters.

## Notes

### Competing Interest Statement

The authors have declared no competing interest.

### Summary of Updates

Substantially updated and revised, notably to include new RNAseq data in Figures 2, 3, and 8 as well as additional experimental work in Figure 6. Text has been restructured and revised.

